# DeepLocRNA: An Interpretable Deep Learning Model for Predicting RNA Subcellular Localization with domain-specific transfer-learning

**DOI:** 10.1101/2023.11.17.567519

**Authors:** Jun Wang, Marc Horlacher, Lixin Cheng, Ole Winther

## Abstract

Accurate prediction of RNA subcellular localization plays an important role in understanding cellular processes and functions. Although post-transcriptional processes are governed by trans-acting RNA-binding proteins (RBPs) through interaction with cis-regulatory RNA motifs, current methods do not incorporate RBP-binding information. In this paper, we propose DeepLocRNA, an interpretable deep-learning model that leverages a pre-trained multi-task RBP-binding prediction model to predict the subcellular localisation of RNA molecules via fine-tuning. We constructed DeepLocRNA using a comprehensive dataset with variant RNA types and evaluated it on held-out RNA species. Our model achieved state-of-the-art performance in predicting RNA subcellular localization in mRNA and miRNA. It has demonstrated great generalization capabilities, not only for human RNA but also for mice. Moreover, the interpretability of the model is enhanced through the motif analysis, enabling the understanding of the signal factors that contribute to the predictions. The proposed model provides general and powerful prediction abilities for different RNA and species, offering valuable insights into the localisation patterns of RNA molecules and contributing to advancing our understanding of cellular processes at the molecular level.

## INTRODUCTION

RNA localisation is the process of transporting and anchoring RNA molecules to specific subcellular regions, where they can perform their functions in gene expression, cell differentiation, and development^1–4^. The misregulation and perturbation of RNA localization are relevant to various disease phenotypes, including cancer^5–8^, development disorders^9–12^, and disorders involving neuromuscular or neuronal dysfunction^13–17^. To play a role in cellular regulation, RNA molecules are transported from the nucleus to target compartments and regulated by RNA binding proteins through three primary mechanisms: 1) direct transport, 2) protection from mRNA degradation, and 3) diffusion and local entrapment^2^. All of these localization mechanisms require coupled protein components to interact with the RNAs to form a ribonucleoprotein (RNP) complex. This essential interaction is primarily driven by *cis-*regulatory elements, also known as zip codes, which serve as key factors in the linear RNA sequence or structure. They determine the interaction between RNA and the RNA binding domain (RBD) of RNA binding protein (RBP)^18^, directing RNA to designated organelles.

Characterizing the factors involved in a RNP complex is important for understanding how RNA traffics from its nascent state in the nucleus to regions outside the nucleus. Cross-linking and immunoprecipitation followed by sequencing (CLIP-seq) is the most common protein-centric experimental approach to measure the protein-RNA interaction profile across the whole transcriptome. Specifically, the method employs UV light to create an irreversible covalent bond between proteins and RNA in their immediate vicinity. This is done before immunoprecipitation purification, protein digestion, cDNA library sequencing and bioinformatics analysis^18^. There are several variants of CLIP, such as individual-nucleotide resolution CLIP (iCLIP)^19^, enhanced CLIP (eCLIP)^20^, and m6A individual-nucleotide resolution UV crosslinking and immunoprecipitation (miCLIP)^21^, which have different modifications in their purification and cDNA library preparation, enabling the protection of the truncations in the protein-RNA interaction sites that helps to increase the specificity and reach to the single nucleotide resolution of the RNA-protein interaction detection.

Currently, there are several machine learning-based tools available for predicting the localization of transcripts. These tools can be broadly categorised into two main types - image-based and sequence-based models. Image-based models leverage manually curated features to characterize RNA distributions^22,23^ or employ cutting-edge computer vision methods to learn hidden feature representation^24^. Sequence-based models^22,23,25^ (see review^4^) predict the localization derived from the primary sequence. The inherent features of *cis-*regulatory elements and the secondary structure are biologically relevant for determining where transcripts should be transported through binding with RNA-binding proteins (RBPs). However, predicting localization exclusively based on the primary sequence may have inherent defects as the primary sequences themselves do not contain RBP binding information. A single sequence can bind with different RBPs, indicating that the regulation of RNA trafficking is a sophisticated and systematic RNA-protein binding network. Ideally, measuring transcriptome-wide RNA-protein interactions would deliver a broad interaction profile between RBPs and RNAs, revealing the numerous regulatory aspects of co-and post-transcriptional gene expression, including RNA splicing, polyadenylation, capping, modification, export, localization, translation and turnover^24,26^.

In this study, we propose DeepLocRNA, an RNA localization prediction tool based on fine-tuning of a multi-task RBP-binding prediction method, which was trained to predict the signal of a large cohort of eCLIP data at single nucleotide resolution. We demonstrate that our model can gain performance from the learned RBP binding information to downstream localization prediction across 4 RNA species, and perform robustly to predict the localization with a limited training dataset. Furthermore, we also apply our model on training in multiple species data and extend the application in a biologically interpretable manner. A user-friendly web server is available at: https://biolib.com/KU/DeepLocRNA/

## MATERIAL AND METHODS

### Localization data source

The localization data used in this study were initially collected from the RNALocate2.0 database^27^ (http://www.rna-society.org/rnalocate1/), which provides the annotated RNA localization information supported by experimental evidence. Then, we retrieve the paired RNA sequences from different sources. For the mRNA dataset, we simply include the mRNA benchmark dataset from DM3Loc^25^. When collecting the human and mouse datasets, we employ the following process and data source to get the sequence. For snRNA, snoRNA, and lncRNA, we downloaded most of the sequences from the NCBI RefSeq database (https://ftp.ncbi.nlm.nih.gov/refseq/H_sapiens/annotation/GRCh38_latest/refseq_identifiers/G <u>RCh38_latest_rna.fna.gz</u>). Alternatively, sequences were extracted from the Ensemble database (https://ftp.ensembl.org/pub/current_fasta/homo_sapiens/ncrna/Homo_s <u>apiens.GRCh38.ncrna.fa.gz</u>), and RNAcentral (https://ftp.ebi.ac.uk/pub/databases/RNAcentr <u>al/current_release/sequences/rnacentral_species_specific_ids.fasta.gz</u>) if they were not present in NCBI. The miRNA sequence was downloaded from the miRBase (https://www.mirbase.org/). To remove the confusion of the gene names and their synonyms in the dataset, we downloaded the FTP file (https://ftp.ncbi.nlm.nih.gov/refseq/H_sapiens/Ho <u>mo_sapiens.gene_info.gz</u>) to match the unique NCBI id and gene symbol for each gene when available in the NCBI. Alternative splicing selectively includes or excludes exons while RNA processing, resulting in variant transcripts of a certain gene. Here, we select the longest transcripts if multiple transcripts are retrieved.

### Unified benchmarking dataset

After getting all sequences that align completely with the localization label provided by RNALocate 2.0, we built a comprehensive dataset for humans and mice, including mRNA, lncRNA, miRNA, snRNA and snoRNA To specify the target, we selected and eliminated redundant representations, assigning the labels "Nucleus," "Chromatin," "Nucleoplasm," "Nucleolus," and "Nuclear" with the same target label as "Nucleus". Similar to DM3Loc^25^, we also kept cytoplasm to assist training by data augmentation, while clearing it in the testing step. The single-label genes were mixed with multilabel genes to form a rich data source for learning the representative localization patterns. Consequently, the pooled 8 most abundant target compartments were kept. To prevent data leakage, we employed CD-HIT-EST^28^ to eliminate redundant sequences. We employed a two-step process to eliminate redundant sequences when pooling all sequences. In the first step, we filtered the intra-dataset by establishing selection criteria, which included a similarity threshold of 0.9 for lncRNA and 0.95 for miRNA, snRNA and snoRNA. These criteria considered factors such as abundance and sequence length. Sequences of mRNA were preprocessed to be non-redundant, as described in a prior work^25^, with a cutoff of 80% sequence similarity. In the second step, we amalgamated all the sequences and set a similarity threshold of 0.95 to retain the majority of the short sequences. This two-step approach is designed to preserve the majority of short sequences (<100nt), as it filters out the most similar long sequences in the first step while allowing individual short sequences to form distinct clusters in the second step.

This process resulted in the following number of remaining genes within specific compartments: Nucleus (13,352), Exosome (22,335), Cytosol (2,587), Cytoplasm (10,026), Ribosome (5,226), Membrane (3,356), ER (1,977), Microvesicle (1,958), and Mitochondrion (33) (Supplementary Table 4), and the mRNA takes the majority part of the final filtered genes in each compartment (Supplementary Figure 5). This comprehensive pooled benchmarking dataset was used to train a unified model in which 5 RNA species were involved across 8 compartments. Curated datasets were split into 5-fold subsets according to the RNA types and the distribution of the constitution of localization. For example, genes with labels as “111000000”, which means they have the label of Nucleus, Exosome, Cytosol, will be split accordingly in mRNA and miRNA if they exist in these two RNA species. Otherwise, only one of them will take each fold.

### Independent benchmarking dataset

To assess the performance of various methods applied to a specific RNA type, we curate a subset of RNA data from our unified benchmark dataset for rigorous benchmarking analysis. This benchmark dataset exclusively comprises RNA localization data sourced from Homo sapiens, encompassing 7 distinct cellular compartments in mRNA, namely the Nucleus, Exosome, Cytosol, Cytoplasm, Ribosome, Membrane, and ER. In the case of lncRNA, we focus on 5 compartments: Nucleus, Exosome, Cytosol, Cytoplasm, and Membrane. miRNA was labelled as extracellular and intracellular to match the comparison with iloc-miRNA^29^. snoRNA are not subjected to division due to the absence of predictive tools for comparison.

Our independent dataset primarily comprises data from three RNA types: mRNA, lncRNA, and miRNA. mRNA data exhibit 58 distinct label combinations, ranging from single label to a maximum of seven labels. The Exosome label stands out as the most popular label, and is nearly twice as prevalent as the next most-popular label, which is a combination of both nucleus, and cytoplasm. Similarly, in lncRNA, we observe 21 label combinations, yet none of the lncRNAs are associated with the ER and membrane compartments. However, we also found 21 combinations in miRNA which are also dominanted by a combination of exosome and microvesicle (Supplementary Figure 3).

### Mouse dataset

Mouse sequence data were processed the same as it was implemented in the human unified dataset, including reducing the redundant sequences and train test split. After carefully selecting the curated mouse data, we get the integrated dataset with three RNA types, lncRNA, miRNA, and mRNA, across 4 variant compartments, Nucleus 2271, Exosome 1116, Cytoplasm 1520, Mitochondrion 96 (Supplementary Table 4). Here we kept the cytoplasm because the cytosol compartment only has dozens of genes that are filtered out in our selection stage. All data were split into 5-fold and used for cross-validation. No tools are available for predicting mouse RNA localization, so we didn’t split the mouse unified benchmarking dataset into subsets to compare different methods.

### Model structure

The architecture of DeepLocRNA is provided in Supplementary Figure 6. It is an end-to-end differentiable model that consists of a pre-trained RBP sequence-to-signal encoder, followed by an attention block and ending in a multi-class classification head. RBP binding signals are extracted to supervise the CNN to focus not only on the sequence composition but also on the RBP potential binding signals. After the RBP-aware encoding, a self-attention layer is applied to allow the model to extract information from relevant parts of the sequence. The attention layer maps from sequence to a fixed length representation that is then fed into a simple fully connected classification network.

### RBP backbone model

Transcript localization is largely governed by an intricate interplay of RNA-binding proteins (RBPs)^4^. Consequently, RBP-binding information is expected to boost the accuracy of localization predictions. To obtain the RBP binding signal, derived from the interaction between RNAs and RBPs that principally guides the RNA trafficking to the target compartments, a pre-trained RBP model was built. We selected 8000 nt as our input length. For sequences longer than 8000 nt, we truncated both ends to keep both 5’ and 3’ information. Sequences shorter than 8000 nt were padded to reach this length. As a result, all sequences were standardized to 8000 nt, ensuring a consistent input length. We take an 8000 nt RNA sequence as input, which is one-hot encoded by mapping the bases A, C, G and U to binary vectors [1, 0, 0, 0], [0, 1, 0, 0], [0, 0, 1, 0] and [0, 0, 0, 1], respectively. The architecture of our RBP backbone model was adapted from RBPNet^30^, and the architecture is as follows. First, an initial 1D convolution layer with 512 filters of size 7 acts as a motif extract and projects the RNA sequence into a high-dimensional space. This is followed by 10 residual blocks, each containing a 1D convolution operation with filters of size 4, batch normalization, ReLU activation and dropout with a probability of 0.3. The first 6 residual blocks contain 384 filters, while the last 4 blocks contain 512 filters. Later, the output of the last residual block then serves as input to 256 output heads (Supplementary Figure 6). A dilation factor of 1.5^i is applied to the convolution operation of each block, where i = {0, …, 9} is the depth of the residual block. Finally, a point-wise convolution operation with 223 filters and linear activation projects the feature map of the last residual block to logits parametrizing a multinomial distribution of eCLIP crosslink counts, in analogy to RBPNet^30^. The loss was computed as the sum of negative log-likelihoods across the 223 eCLIP tracks. For training set construction, the human transcriptome CRCh38.p13 was first tiled into fixed-length 1000nt regions using a sliding-window approach with a stride of 750. For each region, the corresponding RNA sequence together with count-vectors for each eCLIP track were extracted. Samples were then split chromosome-wise into train, validation and test sets, following the split of RBPNet^30^ and the model was trained for 50 epochs using the Adam optimizer with a learning rate of 0.001. When doing fune-tuning, a pooling layer as size of 8 was introduced, leading to an effective sequence length of 1000.

### Attention block

To enhance the model’s focus on the functional regions of the sequence, DeepLocRNA incorporates an attention mechanism, a widely utilized technique in various fields, including document classification^31^. The attention mechanism comprises three fundamental components:

The score function calculates attention scores and is defined as follows:

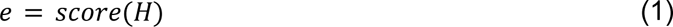

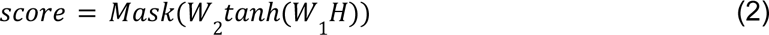

In this scenario, we suppose the attention score function takes the output of the backbone model as the input matrix *H* ∈ R^256xT^, where 256 is the final embedding dimension of the backbone model and *T* is the sequence length after pooling. To get the attention score *e*, the score function was introduced, where *W*_1_ ∈ R^ax256^ serves as a weight matrix. *W*_2_ ∈ R^hxa^ defines the head of the attention mechanism, where h represents the attention heads. To introduce non-linearity and facilitate gradient propagation, the *Tanh* function is employed. Furthermore, the mask function *M* plays a crucial role by imposing a substantial penalty of -10000 on the padded sequence, effectively diverting the attention away from these regions.

Next, the attention scores are further processed by the alignment layer to align the attention score within [0, 1] via the alignment function.

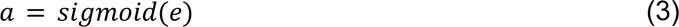

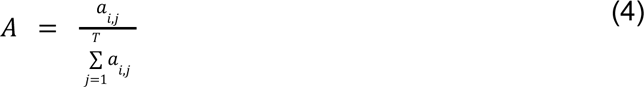

Where *a_i,j_* is the attention weight after alignment by sigmoid function. To make the attention results more smoother, we introduce a softmax transformation to transfer the attention weight into a probability distribution A across the attention heads.

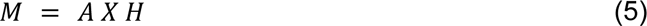

Finally, the transformed attention weights are multiplied by the input matrix to get the context value as the output of the overall attention block.

### Model training

In order to minimize the difference between true multilabel and predicted probabilities, we employed a binary cross-entropy loss function tailored for multilabel classification tasks. When benchmarking the mRNA model with DM3Loc, we employed weight binary cross-entropy. Because of the label inconsistency, we exempt the weights to calculate the loss function while training the unified model.

As the data from each class have clear imbalance issues, we took class weights into account to address data imbalance challenges. The weight scheme in the loss function was formulated as follows:

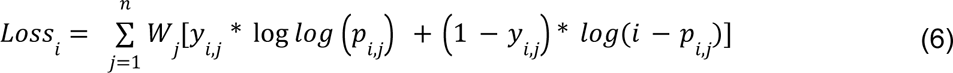

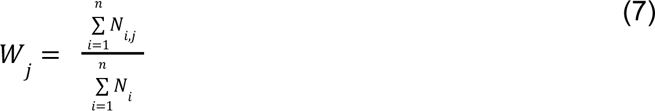

where *y*_*i*_ ∈ {0, 1} is the true label and *p*_*i*_ ∈ [0, 1] denote the predicted probability values of the model. There are variant labels in different training schemes. For example, we utilized 7 compartment labels while training the mRNA model. Hence, each *y*_*j*_ indicates 7 labels in n ∈ {1,2, …, 7}. Whereas, in the case of the miRNA model, *y*_*j*_ can take 5 different labels from n ∈ {1,2, …, 5}. Furthermore, the weight of each class *W*_*j*_ was defined by the proportion of each class accordingly. The *Loss*_*i*_ was summed up by all samples before getting the average Loss in each batch.

To stabilize the training process and prevent gradient-related challenges, we applied a gradient clip during backpropagation. Specifically, the gradient was stored as a vector and the L2 norm of the gradient was calculated.

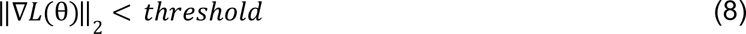

We check whether the L2 norm of the gradients exceeds a predefined threshold (here we set as 1) (8). If the L2 norm of the gradient ||▽*L*(θ)||_2_ exceeds the thresholds we set, we rescale the gradient as follows:

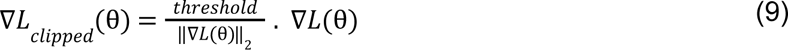

This step ensures that the magnitude of the gradients was under control, preventing any exploding gradient issues while training. We also used the Adam stochastic optimization method with a learning rate of 0.005 and set the weight decay as 1e-5 to prevent overfitting in the way of L2 regularization.

When training the unified model with multiple RNA species, we incorporate identity tags with four dimensions to represent each RNA type within the fully connected layer preceding the classification heads. These tags enable the model to discern distinct inputs, enhancing its capacity to extract varied mechanisms governing the localization of different RNA species and enable multi-RNA prediction.

The entire model was trained based on the PyTorch deep learning framework, and PyTorch-lightning, a lightweight PyTorch wrapper, was implemented to simplify the process of organizing and training the PyTorch model. PyTorch-lightning streamlined the training workflows, automating tedious tasks such as setting up training loops, handling device placement on 4 NVIDIA A100 GPUs with 40GB memory underlining the DDP (Distributed Data Parallel) strategy, which keeps repeats of the model in different GPUs and split the data while training synchronously. This not only saved valuable development time but also ensured the efficient utilization of powerful hardware resources.

### Model evaluation

In our model evaluation, we employed a comprehensive assessment approach, focusing on four key performance metrics: F1 score, Matthews Correlation Coefficient (MCC), Area Under the Receiver Operating Characteristic curve (AUROC), and Area Under the Precision-Recall curve (AUPRC). The AUROC and AUPRC were specifically utilized to gauge the model’s robustness, and the F1 score and MCC were employed to evaluate the model’s statistical accuracy.

To determine these optimal thresholds when using MCC and F1 scores, we leveraged the test dataset to identify the threshold for each RNA compartment that yielded the highest MCC. Separate thresholds were established for various RNA types, with these final thresholds subsequently applied to the predictive server (Supplementary Figures 8-12) For instance, in the context of mRNA classification, 0.7551 for the nucleus, 0.9796 for exosome, 0.2245 for cytosol, 0.2857 for ribosome, 0.3061 for membrane, and 0.1837 for the ER.

### Model explanation

We consider approaches to model explanation: built-in attention weights and the Integrated Gradient (IG)^32^ attribution method.

To provide a clearer illustration of the attention mechanism, the attention weights serve to showcase how the model dynamically directs its focus onto the sequence. Given that the majority of *cis-*regulatory elements were predominantly found at two ends of the sequence, we selectively truncated the sequence to keep these critical regions. Specifically, we focused our analysis on mRNA sequences to maintain both the 5’UTR and the 3’UTR. Sequences exceeding 2000 nt were selected, and 2000 nt were trimmed from two ends to establish a uniform sequence length. For the computation of attention weights, we calculated z-scores across attention heads and determined a mean value over the pooled sequence length of 1000. Subsequently, we applied min-max normalization to standardize the attention weights within a range of 0 to 1 for enhancing visualization. To restore the full length we simply replicated the pooled sequences 8 times to get back to 8000 nts.

We used Integrated Gradients (IG)^32^ to extract critical motifs with a high level of informativeness, essential for RNA localization prediction. To enhance our analysis, we divided the dataset into eight distinct compartments, allowing us to pinpoint the most frequently occurring and influential motifs within each compartment. The overall IG scores were computed using sequences truncated to 2500 nucleotides from both the 5’ and 3’ ends. Subsequently, we aggregated attribution scores for each position within the sequence across four nucleotide dimensions. We identified 5-mer motifs by sliding a 5-nucleotide window across the 8000-sequence length, selecting the 5-mer with the highest IG score for each sequence. Next, we pinpointed the top 5 maximum attribution values within each compartment dataset, representing the most impactful motifs driving sequence trafficking. Finally, these top 5 effective motifs for each compartment were compared with the top 2 motifs extracted from the RBPnet dataset^30^.

### Benchmarking the models

During the benchmarking of mRNA predictive tools, there was a potential overlap between the training data used for iLoc_mRNA, mRNALoc and our benchmark test dataset. The data leakage could artificially improve model performance and cause the benchmark analysis to become unfair. Therefore, we excluded the genes in our benchmark test that were involved in iLoc_mRNA and mRNALoc training step while doing benchmark comparison.

Specifically, to mitigate this issue, we filtered out approximately 900 genes from each fold of the test data used for iLoc_mRNA. Additionally, when calculating performance metrics, we excluded multiple label predictions generated by iLoc_mRNA and instead focused on four unique labels - Nucleus, Cytosol, Ribosome, and ER - for binary evaluation while setting the cutoff as 0.5. To evaluate mRNALoc, we accessed the webserver and downloaded the standalone tool from http://proteininformatics.org/mkumar/mrnaloc/download.html. We removed any benchmark test dataset overlapping with their training data. Any predictions labeled as “Extracellular_region”, “Mitochondria”, and “No Location Found” were assigned a value of “0” to differentiate them from the true label. Predictions within our benchmark dataset labels were assigned their predicted values. We set a cutoff of 0.1 for mRNALoc when evaluating using the benchmark test dataset. The standalone version of DM3Loc was downloaded from https://github.com/duolinwang/DM3Loc. To parallel evaluate DM3Loc, we retrained the model in 5-fold cross-validation with the same non-redundant multilabel benchmark dataset.

In the evaluation of lncRNA predictive performance, we employed a suite of tools, including LncLocator (http://www.csbio.sjtu.edu.cn/bioinf/lncLocator/), DeepLncLoc (https://github.com/ <u>CSUBioGroup/</u>DeepLncLoc), and iLoc-lncRNA (http://lin-group.cn/server/iLoc-LncRNA/predi ctor.php). In cases where downloading the standalone tools proved unfeasible, we resorted to utilizing their respective web servers to conduct the predictions.

For the benchmarking of miRNA, we relied on the web server predictions of iloc-miRNA. To facilitate the evaluation process, we categorized all compartments into two overarching groups: extracellular and intracellular, and computed the relevant metrics separately for these distinct partitions.

### DeepLocRNA webserver

We provide a user-friendly web server, https://biolib.com/KU/DeepLocRNA/, powered by the Biolib library that has been developed to provide secure access to bioinformatics tools directly within the browser. Users can obtain predicted localization results by uploading a FASTA-formatted file or downloading the locally installable version of DeepLocRNA. The server supports optional specification of species and RNA types for running the prediction model.

## Results

### Model construction

The workflow of our model construction begins with a backbone model that is pre-trained to predict RBP-binding profiles for a set of eCLIP datasets from the ENCODE database^33^, followed by an attention layer and a fully connected layer, before the classification heads (Figure 1). In the initial pre-training stage, the model is encouraged to detect RBP binding sites, thereby learning a rich representation of an RNA sequence, conditioned on its trans-acting factors (Methods). We hypothesize that the pre-trained hidden representations encapsulate the broadly underlying principles governing RBP-RNA interactions, thereby learning a rich representation of an RNA sequence, conditioned on its trans-acting factors. Potentially, a subset of these interactions plays a crucial role in guiding RNA localizations.

**Figure 1.**
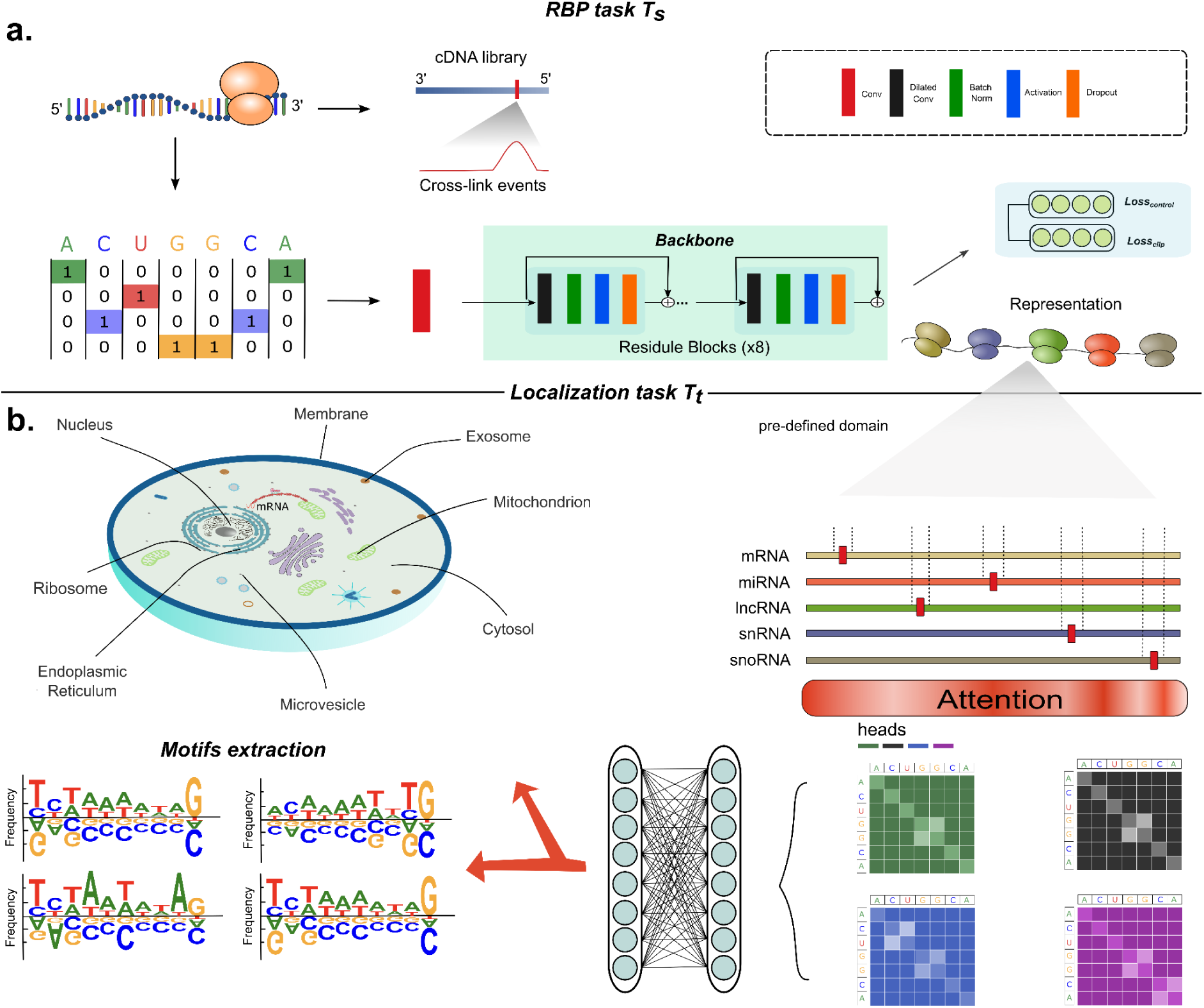
A comprehensive visualization of the pre-training and fine-tuning schemes used in localization prediction. **a.** Sequences are one-hot encoded before serving as the input to our RBP sequence-to-signal model, which enables the prediction of the RBP binding signal in a single nucleotide resolution. After eight rounds of feature extraction, the final feature embeddings were generated to yield the representation of protein-RNA interaction, getting the **T_s_** pre-trained model. **b**. RBP binding signals are used to guide the localization prediction across 8 compartments **T_t_**. Before going to the fully connected layer, the self-attention mechanism is used to try to attend to the biological functional zip code motifs. When the localization results are predicted, functional motifs can be extracted to do the model interpretation derived from the IG score across 4 nucleotide dimensions.

To achieve this objective, we undertook a series of technical steps to modify the backbone model. Firstly, we removed the output heads from the backbone model to extract critical RBP binding information. As a result, the model generated embedding vectors of the shape [8000,256]. Subsequently, an attention layer was strategically incorporated to capture the salient regions within the sequences. Additionally, a fully connected layer was added to further extract the feature, along with new classification heads to facilitate RNA localization prediction (Figure 1).

DeepLocRNA was subsequently fine-tuned with a rich diversity of RNA localization data, encompassing long non-coding RNA (lncRNA), micro RNA (miRNA), small nuclear RNA (snRNA), small nucleolar RNA (snoRNA), and messenger RNA (mRNA) for multi-RNA localization prediction (Supplementary Table 4). During the fine-tuning of the model, we observed optimal performance when half of the intermediate layers were unfrozen, allowing the model to learn localized and contextually relevant RBP binding representations (Supplementary Figure 1).

### Benchmarking with the other tools

We divided the mRNA dataset from our benchmarking dataset to ensure a fair comparison with three other predictive tools: DM3Loc^25^, iLocmRNA^35^, and mRNALoc^22^. In this evaluative process, to ensure a fair comparison among different methodologies on the same test data, we carefully filtered out genes in the testing dataset of each fold that had been previously used in the training process of iLoc-mRNA and mRNALoc, thus mitigating the risk of data leakage. As different tools may exhibit uneven target labels across cellular compartments, our evaluation focused solely on testing within the available compartments of each tool. For example, since mRNALoc was constructed only for the Nucleus, Cytoplasm, and Endoplasmic Reticulum (ER), we exclusively computed metrics for these supported compartments. The results unequivocally demonstrate that fine-tuned DeepLocRNA outperforms the other three methods in terms of overall performance across five compartments with a high macro AUROC of 0.7493. When training the model from scratch, which means we reset all the parameters, a lower AUROC of 0.7283 was obtained (Table 1). In comparison to DM3Loc, the previous state-of-the-art mRNA localization predictor, our model exhibits considerable advancements in Exosome localization prediction (AUROC from 0.7273 to 0.7633, Supplementary Table 1). Although its predictive capability for the ER does not match that of DM3Loc, we achieved higher performance by assigning weights to the loss function based on the abundance of each compartment (Supplementary Table 1). The training strategy employing early stopping also illustrates a more rapid descent in loss and a lower final loss value when compared with DM3Loc (Supplementary Figure 2).

**Table 1.** The average performance of DeepLocRNA across four RNA species. The number in bold represents the max value across different tools. The macro average represents the mean value of specific metrics across five different compartments.

| RNA types | Tools | MACRO-F1 | MACRO-MCC | MACRO-AUROC | MACRO-AUPRC |
| --- | --- | --- | --- | --- | --- |
| mRNA | DM3Loc | 0.4315 | 0.1713 | 0.7423 | 0.5743 |
|  | iLoc-mRNA | 0.1832 | 0.0441 | 0.5248 | 0.3093 |
|  | mRNALoc | 0.3441 | 0.0497 | 0.5211 | 0.4283 |
|  | DeepLocRNA-ind<br>(training from scratch) | 0.3191 | 0.0643 | 0.7283 | 0.5621 |
|  | DeepLocRNA-ind<br>(instructive fine-tuning) | **0.4647** | **0.1774** | **0.7493** | **0.5786** |
|  | DeepLocRNA-uni<br>(instructive fine-tuning) | 0.4075 | 0.158 | 0.7433 | 0.5706 |
| miRNA | DeepLocRNA-uni<br>(training from scratch) | 0.5446 | **0.4282** | 0.8671 | **0.6809** |
|  | DeepLocRNA-uni<br>(instructive fine-tuning) | **0.5683** | 0.418 | **0.8682** | 0.6769 |
| snoRNA | DeepLocRNA-uni<br>(training from scratch) | 0.5037 | 0.0109 | 0.5843 | 0.1972 |
|  | DeepLocRNA-uni<br>(instructive fine-tuning) | **0.5293** | **0.0248** | **0.6452** | **0.2599** |

ind: training from the independent dataset; uni: training from the unified dataset.

Subsequently, we conducted an evaluation of miRNA localization using a dedicated miRNA training dataset. In the past, the majority of research efforts have been directed toward developing models for mRNA and lncRNA localization prediction^4^. We only found iLoc-miRNA available for predicting miRNA trafficking, which offers predictions primarily distinguishing between extracellular and intracellular localization^29^. In our evaluation, we segregated our miRNA dataset into intracellular and extracellular segments for a comprehensive and equitable comparison with iLoc-miRNA. DeepLocRNA consistently demonstrates the beneficial contributions of pre-trained protein information, outperforming both training-from-scratch and iLoc-miRNA (Table 2). Intriguingly, all models exhibit high scores in according metrics, especially all exceeding 0.90 in AUROC (Table 2). This implies the possible existence of specific *cis-*regulatory elements within the primary sequence, facilitating the model’s adaptability to the data.

**Table 2.** Benchmarking DeepLocRNA in the prediction of miRNAs.

| Tools | Test Ratio | Precision | Recall | F1 | AUROC | AUPRC |
| --- | --- | --- | --- | --- | --- | --- |
| iloc-miRNA |  | **0.8916** | 0.8949 | 0.8932 | 0.9286 | 0.8249 |
|  |  | **0.9943** | 0.8603 | 0.9225 | 0.9159 | 0.9746 |
| DeepLocRNA<br>(training from scratch) | Extracellular(693) | 0.8779 | **0.9111** | 0.8940 | 0.9438 | 0.8763 |
|  | Intracellular(172) | 0.9462 | 0.9117 | 0.9283 | 0.9355 | 0.9885 |
| DeepLocRNA<br>(instructive fine-tuning) |  | 0.8850 | 0.9058 | **0.8952** | **0.9547** | **0.8808** |
|  |  | 0.9531 | **0.9143** | **0.9331** | **0.9419** | **0.9905** |

Finally, we evaluate our method against three other lncRNA prediction tools, including DeepLncLoc^34^, LncLocator^36^, and iLoc-lncRNA^37^. While all the compared methods were trained on data with unique labels, filtering out genes with multiple labels, they displayed limited generalizability, with AUROC values ranging from 0.4904 to 0.5192 across all compartments (Table 3). Even when there were overlapping genes between their training data and our benchmarking testing data, these specific tools failed to exhibit robust performance. Notably, our baseline model, trained from scratch, also outperformed these counterparts, substantiating the efficacy of our proposed model structure. Following fine-tuning of our model with pre-trained RBP interaction information, performance gains were observed across all compartments, particularly in the Exosome compartment, where the AUROC increased from 0.5690 to 0.5832. These advancements underscore the beneficial impact of incorporating RBP information on compartment prediction, with the most substantial improvements noted for lncRNA localization to the Exosome. However, the overall performance of lncRNA localization consistently exhibits lower accuracy, despite our tool ranking as the top performer across all compartments. This suggests that reliance solely on primary sequence information may not yield robust predictions for lncRNA localizations, hinting at potential limitations inherent in the track of lncRNA trafficking. This could be influenced by unconsidered factors such as nuclear localization signal (NLS)^38^, nuclear retention signals (NRS)^39^, or secondary structures of the sequence^40^.

**Table 3.** Benchmarking of DeepLocRNA in the prediction of lncRNAs.

| Tools | AUROC | AUPRC | MCC |
| --- | --- | --- | --- |
| DeepLncLoc | 0.5152 | 0.2619 | 0.0266 |
|  | 0.5002 | 0.9736 | 0.0013 |
|  | 0.4931 | 0.0701 | -0.0137 |
|  | 0.5 | 0.0343 | 0 |
| LncLocator | 0.4913 | 0.2534 | -0.0263 |
|  | 0.5044 | 0.9739 | 0.0073 |
|  | 0.4904 | 0.0701 | -0.0119 |
|  | 0.5000 | 0.0343 | 0 |
| iLoc-lncRNA | 0.5106 | 0.2609 | 0.0248 |
|  | 0.4965 | 0.9735 | -0.0028 |
|  | 0.5192 | 0.0734 | 0.0206 |
|  | 0.5000 | 0.0343 | 0 |
| DeepRBPLoc<br>(scratch) | **0.5477** | 0.3000 | 0.0156 |
|  | 0.5690 | 0.9807 | 0 |
|  | 0.5839 | 0.0959 | 0 |
|  | 0.5929 | 0.0579 | 0 |
| DeepRBPLoc | 0.5443 | **0.3027** | **0.0156** |
|  | **0.5832** | **0.9815** | 0 |
|  | **0.5932** | **0.1004** | 0 |
|  | **0.5938** | **0.0658** | 0 |

### A unified model for multi-task learning

Protein-RNA interactions play a fundamental role in directing the localization of diverse RNA species. We trained the model by the unified benchmarking dataset (Methods), leveraging the rich binding signals encoded within our pre-trained model. With a greater number of sequences incorporated, our model was expected to discern crucial features from diverse RNA compositions and encapsulate the entirety of the binding mechanisms into a unified mode.

This training strategy can be characterized as a form of multi-task and multi-label learning, which enables the training of our unified model across eight different cellular compartments spanning five RNA species. Various RNA species were approached as distinct prediction tasks, and our objective was to combine these tasks into a single, unified model. In order to enable the model to distinguish between different tasks, each RNA species was uniquely identified through the introduction of a specialized tag within the fully connected layer (Methods). Notably, there are currently no existing multi-RNA predictive tools available, prompting us to compare our model against a scenario where training is initiated from scratch, thereby underscoring the performance enhancement attributable to RBP binding interactions.

In the context of mRNA localization prediction, our unified model, which initially outperformed DM3Loc in four compartments when trained on an independent mRNA dataset, also exhibited enhanced performance in the Ribosome compartment (Supplementary Table 2). The AUROC of our model in Ribosome lightly outperforms DM3Loc (from 0.7546 to 0.7607, (Supplementary Table 1 and Supplementary Table 2), underscoring the efficacy of our unified training approach. However, the performance of the endoplasmic reticulum (ER) compartment, which exclusively exists within the mRNA dataset in our pooled unified dataset, remains unaffected by data augmentation and is potentially overshadowed by other compartments. Similarly, for miRNA and lncRNA, we already proved DeepLocRNA performs well when trained independently dataset (Table 2 and Table 3). However, lncRNA exhibits relatively lower performance than other RNA types when trained on pooled and independent datasets, suggesting that the localization of lncRNA may not predominantly depend on primary sequence information (Table 1 and Supplementary Table 2). Furthermore, with the adoption of the unified model, we expanded the scope of prediction beyond the generic extracellular and intracellular categories found in iLoc-miRNA, encompassing five more specific compartments—nucleus, exosome, cytoplasm, microvesicle, and mitochondrion. Our model achieved AUROC scores exceeding 0.9 in the first four compartments. However, it struggled to achieve satisfactory performance in the mitochondrion compartment, with AUROC at a low 0.5556 and F1 score at 0, likely attributed to the constraints posed by a limited training sample size of 32. The comparison with training from scratch also proves the performance enhancement using the instructive fine-tuning method (Supplementary Table 2).

Notably, our unified model exhibits exceptional generalization capabilities, particularly when applied to rare datasets, such as small nucleolar RNA (snoRNA), which consist of limited training data and are susceptible to overfitting in training deep neural networks. The AUROC values in the nucleus and cytoplasm compartments are relatively lower, standing at 0.6595 and 0.6071, respectively. Conversely, the performance in the exosome and microvesicle compartments is notably exceptional, reaching a perfect F1 score of 1 in exosome and a high value of 0.9991 in microvesicle. These results underscore a remarkable accuracy in predicting these two compartments, even for the 112 genes in the testing dataset.

These findings emphasize the effectiveness of our pre-training strategy, which empowers the prediction of localization across a wide spectrum of RNA types and provides the most comprehensive localization prediction capability to date.

### Cross-species prediction

RNA-binding proteins (RBPs) occupy pivotal roles in post-transcriptional gene expression regulation by selectively binding to specific RNA sequences or structures. These conserved protein-RNA interactions are often indicative of underlying biological similarities. To evaluate the generalizability of our computational framework across species, we extended our approach to the mouse RNA localization prediction. Here, we leveraged RBP binding knowledge obtained from human data and performed fine-tuning using mouse-specific datasets. Notably, the mouse dataset, although less extensive than its human counterpart, encompasses only three RNA types localized across four distinct compartments.

Our unified model, as previously described, effectively capitalizes on the presence of more abundant data to bolster the prediction accuracy for smaller datasets—a clear advantage when dealing with scenarios such as snoRNA localization in humans. In mice, regarding lncRNA, with a dataset consisting of a limited number of genes (8, 7, and 5 in the Nucleus, Exosome, and Cytoplasm, respectively), our framework exhibits performance levels similar to those observed in the human lncRNA dataset, with AUROC values hovering around 0.5145 to 0.5970 (Table 4). Nevertheless, it is noteworthy that our framework performs relatively well in mRNA localization and excels in coarse-grained classification tasks, particularly in the Nucleus and Cytoplasm (Table 4). Conversely, in the context of miRNA prediction, it encounters challenges in coarse-grained compartment prediction (Nucleus and Cytoplasm) but demonstrates strong performance in the exosome compartment. These observations suggest the potential existence of distinct *cis-*regulatory element regulation mechanisms governing mRNA and miRNA localization in mice.

**Table 4.** Performance of DeepLocRNA in mouse. The bold numbers represent the larger values when compared with the instructive fine-tuning model (left) and training from scratch model (right).

| RNA species | Compartments | F1 | MCC | AUROC | AUPRC |
| --- | --- | --- | --- | --- | --- |
| mRNA | Nucleus | **0.7704** \| 0.7694 | **0.5389** \| 0.5175 | **0.8414** \| 0.8263 | **0.8088** \| 0.7874 |
|  | Exosome | **0.0370** \| 0.0248 | **0.1064** \| 0.0652 | **0.6268** \| 0.5975 | **0.3150** \| 0.2771 |
|  | Cytoplasm | **0.6700** \| 0.5965 | **0.5229** \| 0.4735 | **0.8405** \| 0.8202 | **0.7449** \| 0.7201 |
|  | **Micro** | **0.661** \| 0.6404 | / | **0.9556** \| 0.9516 | **0.7265** \| 0.7004 |
| miRNA | Nucleus | 0 \| 0 | 0 \| 0 | **0.6317** \| 0.6279 | 0.2796 \| 0.3012 |
|  | Exosome | 0.9123 \| 0.9123 | 0 \| 0 | **0.8125** \| 0.7843 | **0.9504** \| 0.9423 |
|  | Mitochondrion | 0 \| 0 | 0 \| 0 | **0.6980** \| 0.6937 | 0.5344 \| 0.5387 |
|  | **Micro** | **0.7378** \| 0.7378 | / | **0.9666** \| 0.9660 | **0.8463** \| 0.8404 |
| lncRNA | Nucleus | 0.3143 \| **0.4603** | 0.1778 \| **0.3172** | 0.5218 \| **0.5694** | 0.4640 \| **0.5221** |
|  | Exosome | 0.0571 \| **0.1056** | 0.1777 \| **0.1548** | 0.5145 \| **0.5984** | 0.3918 \| **0.4611** |
|  | Cytoplasm | 0.3212 \| **0.1030** | 0.1924 \| **0.1500** | 0.5970 \| **0.6083** | **0.4028** \| 0.3930 |
|  | **Micro** | 0.2546 \| **0.2686** | / | 0.8677 \| **0.8854** | 0.3448 \| **0.3850** |
| Overall | Nucleus | 0.7532 \| **0.7539** | **0.5409** \| 0.5233 | **0.8410** \| 0.8287 | **0.7964** \| 0.7767 |
|  | Exosome | **0.4505** \| 0.4464 | 0.4323 \| **0.4256** | **0.7297** \| 0.7113 | **0.5888** \| 0.5723 |
|  | Cytoplasm | **0.6630** \| 0.5886 | **0.5314** \| 0.4802 | **0.8558** \| 0.8375 | **0.7374** \| 0.7126 |
|  | Mitochondrion | 0 \| 0 | 0 \| 0 | **0.9806** \| 0.9804 | 0.5344 \| **0.5387** |
|  | **Micro** | **0.6617** \| 0.6440 | / | **0.9551** \| 0.9517 | **0.7331** \| 0.7117 |

### Model explanation across full-length

To investigate the inner workings of the model, we calculated attention weights in the nearest layer of the backbone encoder. We selectively included sequences exceeding a length of 2000nt, preserving 1000nt from both the 5’ and 3’ ends. Analysis of the attention weights revealed a notable low degree of importance towards the tips of these sequence ends. A decreasing trend from the 5’ end to the 3’ end was observed (Supplementary Figure 7). This outcome slightly diverges from what DM3Loc found, which utilized pooled feature representations as input to the attention layer^25^. In contrast, our input to the attention layer comprises abstract representations of protein-RNA interactions, suggesting a subtle shift towards a higher likelihood of RNA-binding protein (RBP) binding events on the left side.

Integrated Gradients (IG)^32^ significantly enhances model interpretability by revealing key feature attributions linked to prediction targets, improving our understanding of the deep learning model’s decision process. Elevated scores among the four nucleotide bases signify heightened contributions of specific bases to the target compartments, culminating in the formation of a position weight matrix (PWM). We retained 2500nt from both ends of the sequences, resulting in a total sequence length of 5000nt for IG score calculation. Our analysis revealed consistently high attribution levels at two ends of the sequences, underscoring the substantial contributions of both the 3’UTR and 5’UTR to the localization prediction (Figure 2a).

**Figure 2.**
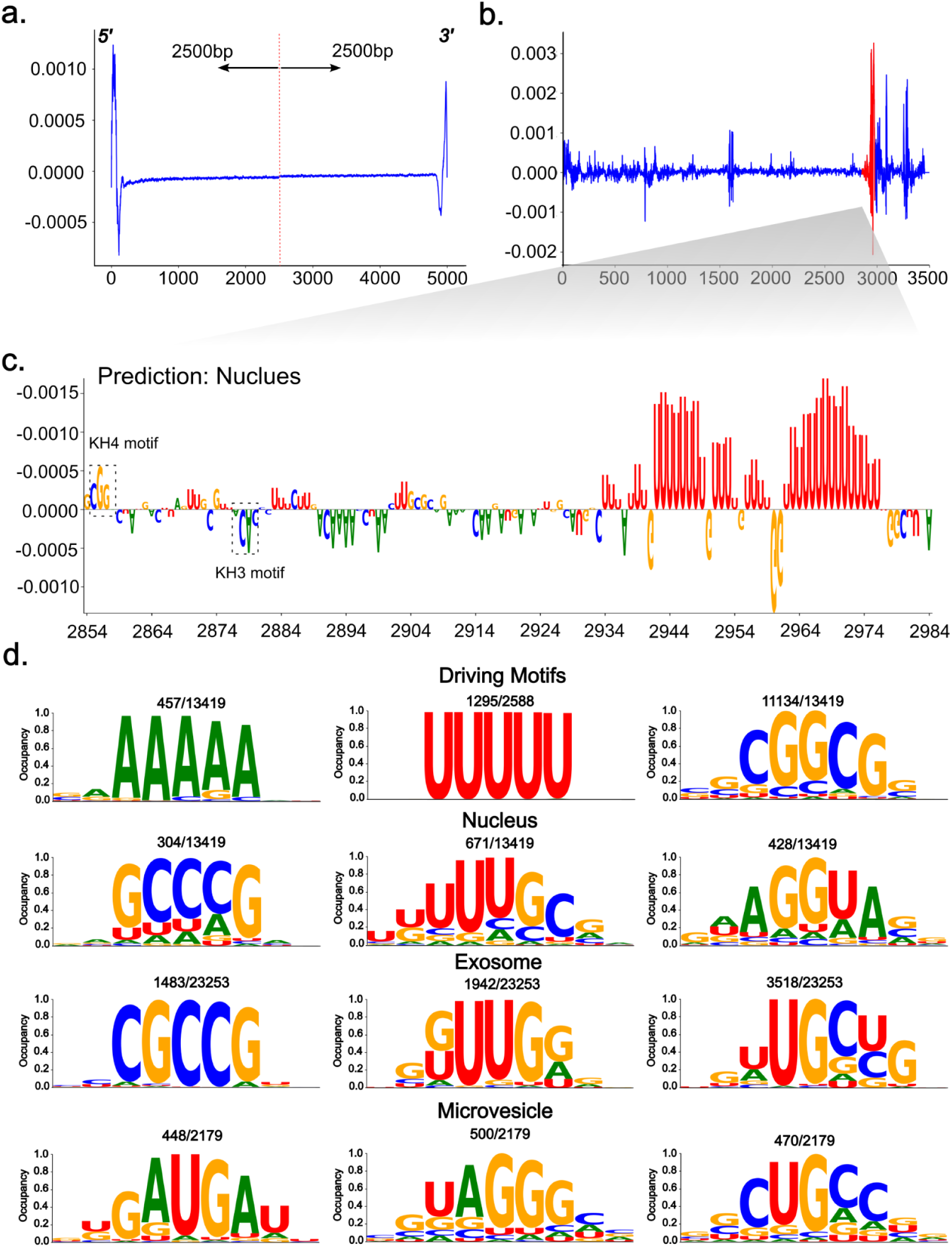
Model explanation with Integrated Gradient (IG) scores and extracted motif visualization. **a**) A visual representation of the attribution score across the two ends of all the sequences in the unified dataset. Sequences exceeding 5000nt have been selected, and 2500nt pairs from each end have been truncated, resulting in a 5000nt sequence length represented on the x-axis. This visualization offers insights into the attribution of importance to different regions at the sequence’s beginning and end. **b**) The IG scores for the ACTB gene. The full length of the gene sequence is displayed, with a red line indicating the *’zip code’* region within the sequence. **c**) A zoomed-in version of the ’zip code’ region from plot b. This plot showcases the attribution score across four dimensions at a single nucleotide resolution. The x-axis commences at the beginning of the *’zip code’* region, allowing for a more detailed examination of the sequence’s key attributes. **d**) The top three 5-mer motifs within the respective localization datasets. The nucleotides displayed in the logo plot represent patterns captured as sliding windows traverse the sequences, and their attribution values are calculated using IG. The mean IG score has been normalized within a range of 0 to 1, as indicated on the y-axis. This analysis unveils crucial sequence motifs and their respective attribution values.

### Model explanation - two case studies

To further validate the biological significance of attribution in target prediction, we downloaded the ACTB gene, which can be translated as the β-actin to form the actin cytoskeleton, from NCBI. The gene’s localization was accurately predicted as the nucleus, demonstrating the robustness of our model’s generalization, especially considering it was not part of our initial benchmark dataset. Subsequently, we computed the IG scores across the full gene sequence and found the highest attribution is localized at 3’UTR of the sequence (Figure 2b). For an in-depth examination of single nucleotide attribution, we manually curated the 52-nucleotide *zipcode* sequence, previously defined as the binding region for RNA-binding protein^41^. Our investigation then focused on the model’s ability to identify motifs associated with ZBP1 and HuD, known to bind overlapping sites within the β-actin zipcode, playing a crucial role in mRNA transport^42^. We observed a positive attribution at the beginning of the *zipcode*, confirming the presence of the KH4 recognition motif 5’- CGGAC - 3’^41^ of the RNA-binding protein ZBP1. Conversely, the KH3 recognition motif 5’- ACAC - 3’^41^ showed a negative attribution (Figure 2c), suggesting a potential structural dependence of ZBP1, making simultaneous binding of KH3 and KH4 unlikely. Both motifs, bound by the ZBP1 protein, are located within the 52-nucleotide region. Notably, U-motifs with the highest attribution scores across the entire ACTB sequence (Figure 2c) were found downstream of this 52-nucleotide region, likely to be bound by HuD, given their preference for U-rich features^43^. When perturbing the 3’ ends with random nucleotides, our model does not predict the nucleus as its compartment, highlighting the robustness of our model in handling perturbation cases.

Huntington’s disease (HD) results from altered HTT gene concentration in the nucleus and cytoplasm, primarily due to expanded CAG repeats^44^. Our model is able to accurately predict HTT localization with variant CAG expansion levels (Supplementary Figure 13a). Specifically, we found that increasing CAG repeats boost the prediction probabilities in both the nucleus and cytosol (Supplementary Figure 13a), with a high level of the mutated CAG attribution (Supplementary Figure 13cde). Removing CAG sequences significantly reduces prediction values (Supplementary Figure 13a). These results underline the importance of CAG in predicting HTT gene localization, which is potentially a valuable target to reduce mutant HTT mRNA accumulation and mitigate the toxic effects of the mutant protein.

### Motif analysis

To investigate whether the modelling of RNA trafficking can unveil inherent functional elements dictating localization predictions in a computational fashion, we compute IG scores across all eight predictable compartments in our model. Given that IG results in our unified model may reflect the collective attribution of multiple RNA-binding proteins, the extracted motifs offer insights into the binding motifs that recurrently appear, possibly due to the dominance of a particular set of highly active RBPs. Thus, we endeavour to ascertain whether these motifs correlate with RBPs sharing similar binding motifs, selecting the top five highly expressive motifs within the respective target set in our unified benchmark dataset.

The identification of the "CGGCG", "A-motif" and "U-motif" motifs emerges as particularly significant driven motifs, successfully classifying five out of eight compartments. Specifically, a high degree of concurrence of the ’U-motif’ is prominently observed in the ACTB zipcode region (Figure 2c), indicative of a compelling binding mechanism that orchestrates the transportation of the gene from the nucleus to the cytoplasm. Furthermore, we unearth motifs specialized for specific localizations. In the nucleus attribution analysis, we discerned that the expressive 5-mer motifs "GCCCG", "AGGUA", and "UUUGC", which are binding motifs of RBMX^43^, KHSRP^45^, TARDBP^46^, regulating alternative splicing localized in the nucleus. Notably, "CGGCG" exhibits a strong correlation with the protein PPRC1, which coactivates nuclear gene transcription (Figure 2d). "CGCCG", found in top motifs of the exosome, is a binding motif identified by FMR1, playing a significant role in endosome cargo loading that often interacts with miRNA^47^. The "GUCCG" element interacts with ZNP2, initially binding to nascent beta-actin transcripts and facilitating binding with ZBP1, associated with nuclear-to-cytoplasmic localization transport^48^. Additionally, motifs not extensively documented in literature yet unique to certain compartments, such as "GUUUC" and "GAUGA," may potentially represent common identification patterns guiding RNA to ER and microvesicle.

Furthermore, we conducted a comparative analysis of these compartment-specific functional motifs with the findings from RBPnet^30^, which predicts the binding interactions between proteins and RNAs. Intriguingly, we identified four distinctive motifs that precisely correspond to the results previously obtained by RBPnet (Supplementary Table 3). Notably, the "U-motif" motif emerged as a prominent motif, featuring prominently among the top two binding motifs for several proteins. For example, FUBP3 has been established as a crucial factor in the regulation of β-actin mRNA, a major constituent controlling RNA mobility and directing its localization through binding to the 3’ UTR^49^. As for the nuclear motif "AGGTA," NCBP2 is intricately involved in various processes, including pre-mRNA splicing, translation regulation, and nonsense-mediated mRNA decay^50^. These intriguing findings warrant further experimental exploration to elucidate the functions of these novel motifs and their interactions with relevant RBPs in the context of RNA localization.

## DISCUSSION

In this study, we address the multi-label RNA localization prediction problem, by leveraging a pre-training scheme to glean protein-RNA binding characteristics at a single nucleotide resolution from the CLIP-seq data. DeepLocRNA thrives when tasked with predicting gene localization based on the guiding influence of RBPs, irrespective of RNA types. Our model also exhibits commendable generalization capabilities in cross-species prediction, particularly in distinguishing between the nucleus and cytoplasm.

Furthermore, we curated a unified, non-redundant benchmark dataset encompassing five RNA types and eight distinct localizations spanning both human and mouse datasets. To enable comparisons with other tools, we dissected the unified dataset, evaluating the performance of our method on subset data. The final model, trained on this comprehensive benchmark dataset, amalgamates sequence information in a data augmentation framework bolstered by pre-trained RBP-RNA interactions. However, the evaluation results indicate that only mRNA and miRNA are feasible in getting favourable results. lncRNA and snoRNA are unable to reach desirable results due to the AUC of these two types still hanging around 0.6.

To analyse predictions using Integrated Gradient (IG), extracting the most informative motifs pertinent to the prediction targets through attribution methods. As a sequence-driven model, DeepLocRNA can be elucidated by examining PWMs across various RNA species, uncovering overarching patterns. In addition to mapping known functional RBP-binding motifs from literature which may represent potential regulatory motifs governing finer organelle localization. These findings hold promise for experimental validation.

Our work represents a pioneering effort in creating comprehensive RNA localization prediction tools employing a sequence-driven approach, blending primary sequence information with RBP binding priors. Future enhancements may involve leveraging large RNA language models, enabling the model to grasp RNA intricacies from genome-wide nucleotide corpora and further refining RNA representation. This adaptable model can also seamlessly integrate diverse data modalities, such as in situ hybridization images, protein expression, and regulation, enhancing its robustness and applicability across various diseases and developmental contexts. Furthermore, we did not account for cell type heterogeneity in our study, primarily because of the requirement for substantial data to train our deep neural network. However, as more data becomes available in the future, it will be imperative to include considerations for cell type heterogeneity in building the model for potential applications. e.g. RNA drug delivery. The wealth of data derived from diverse sources, including microscopy images and RBP binding profiles, paves the way for the development of more precise localization prediction tools, thus facilitating drug discovery and driving novel advancements in disease treatment.

## DATA AVAILABILITY

The unified data for training and testing are available at https://zenodo.org/records/10116380. The standalone tool for local use and source code are available in the GitHub repository, https://github.com/TerminatorJ/DeepLocRNA.

## SUPPLEMENTARY DATA

Supplementary Data are available at NAR online.

## ACKNOWLEDGEMENTS

We like to thank Rachael DeVries who throughly proof-reading this paper.

## Supplementary Information

**Supplementary Figure 1.**
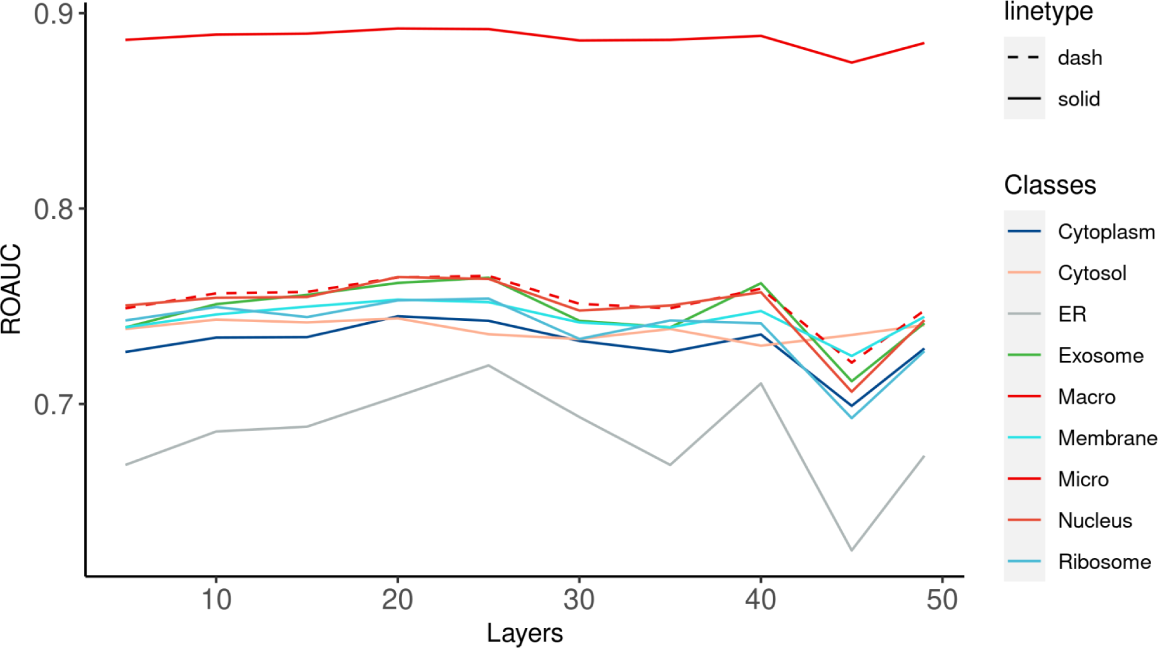
The results of fine-tuning the backbone model. We assess the performance of the DeepLocRNA model in downstream localization prediction by releasing different layers of the backbone model. The x-axis depicts the layers that release from the end of the backbone model to the stem layer, where smaller values indicate higher retention of RBP-RNA binding information. Each compartment is represented by a different colour, except for the two average metrics - macro average and micro average - shown in red. The micro average is visualized as a dashed line, while the macro average is presented as a solid line for distinction.

**Supplementary Figure 2.**
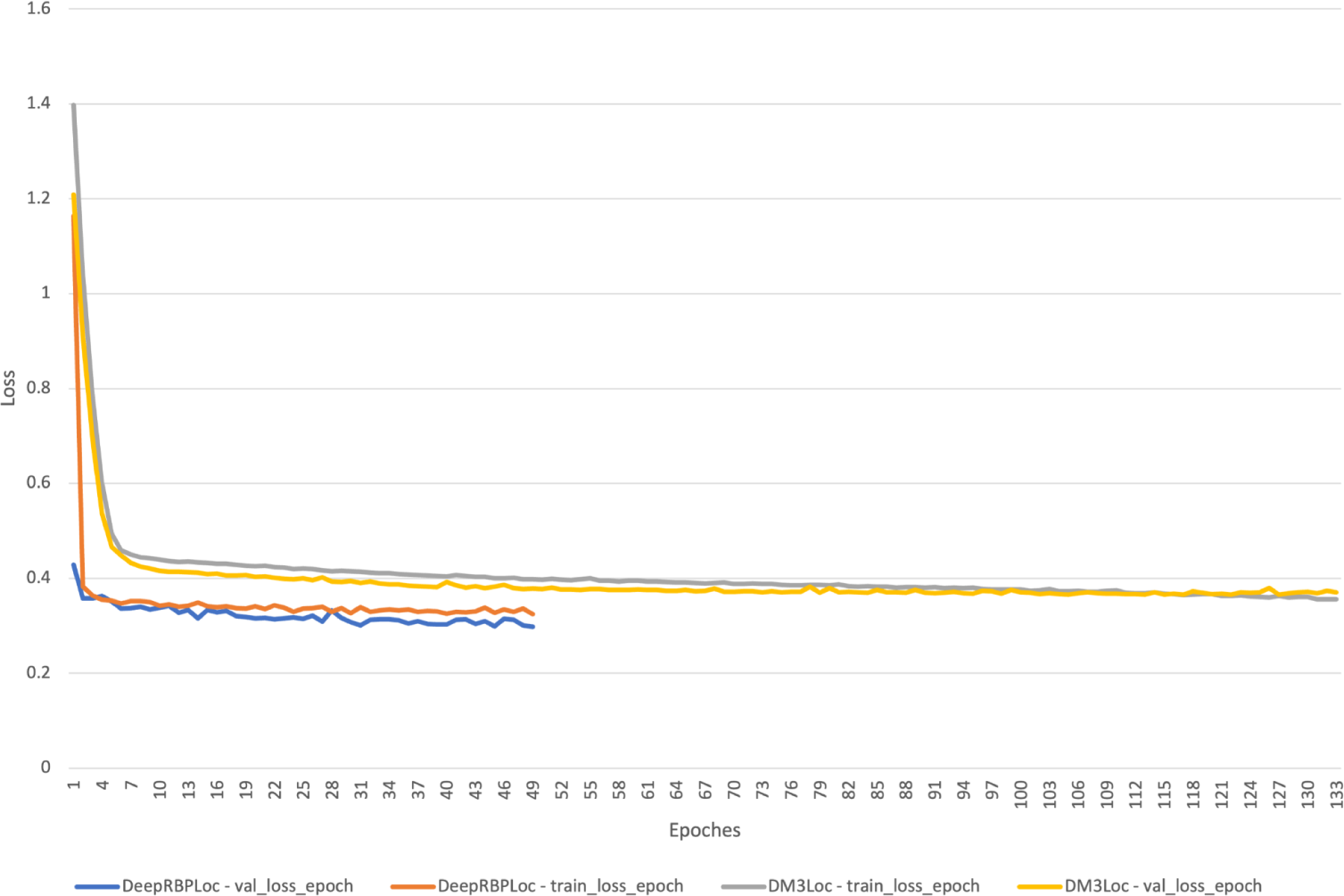
The loss curve during the training of DeepLocRNA and DM3Loc. the x-axis represents the number of epochs needed for model convergence. the y-axis shows the decrease of the binary cross entropy loss while training.

**Supplementary Figure 3.**
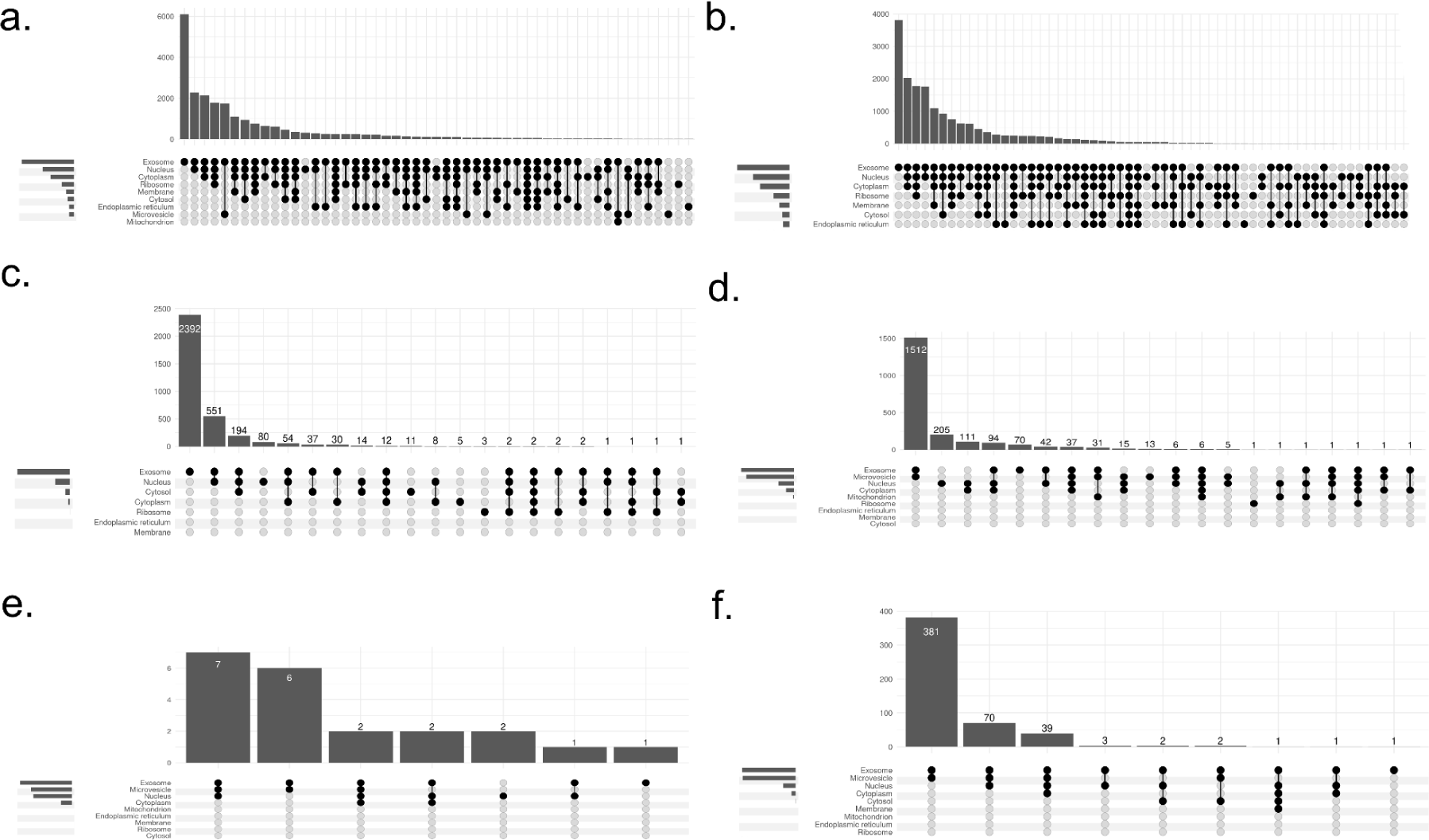
Visualization of the multilabel data. These UpSet plots show different combinations of targets in our dataset. Each column represents a case of multiple labels. The bar in each column indicates its abundance, and the tiny legend on the left side of each plot represents the abundance of each compartment. a) all RNA; b) mRNA; c) lncRNA; d) miRNA; e) snRNA; f) snoRNA.

**Supplementary Figure 4.**
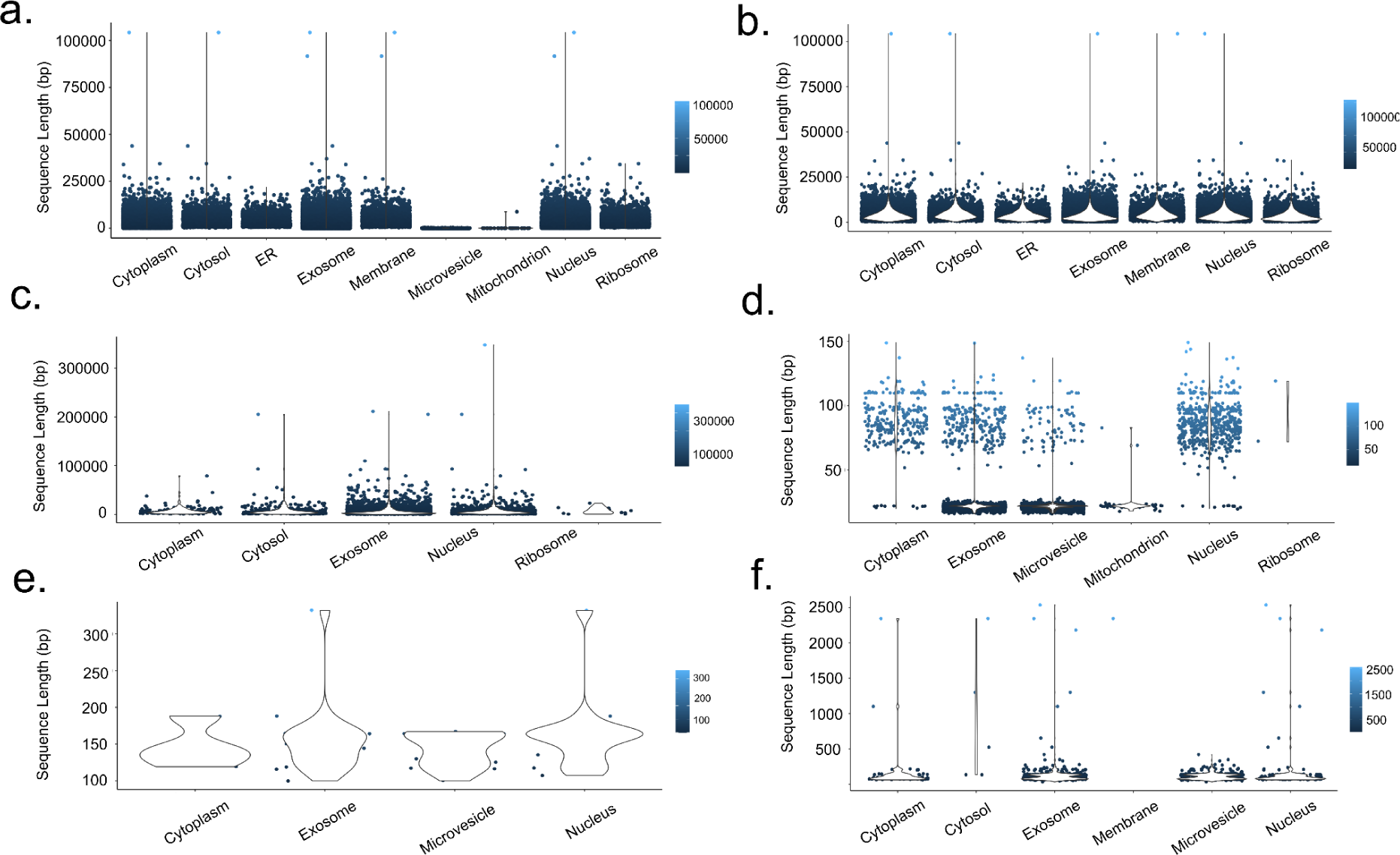
Violin plots of sequence length distribution across all available compartments. a) all RNA; b) mRNA; c) lncRNA; d) miRNA; e) snRNA; f) snoRNA.

**Supplementary Figure 5.**
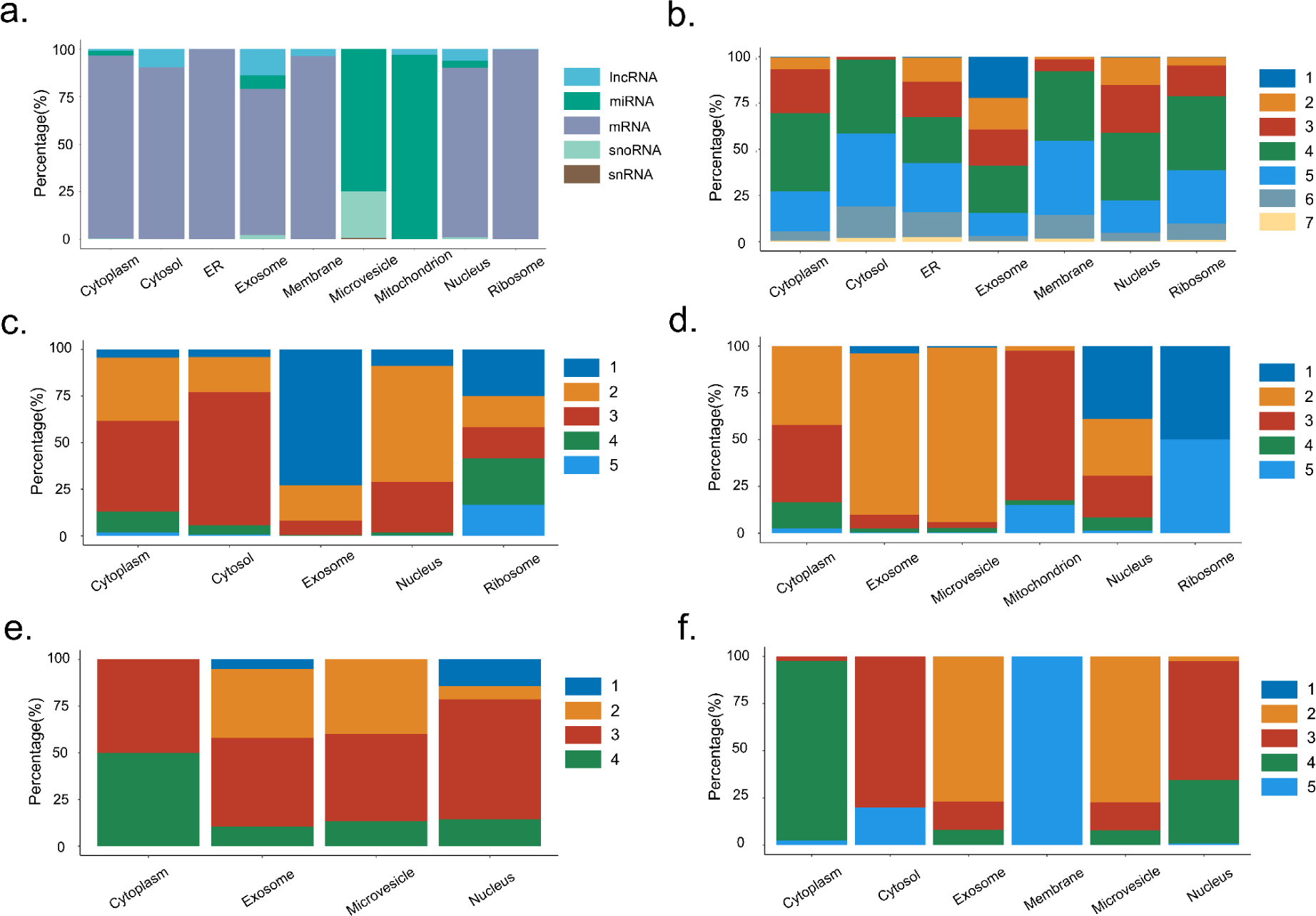
The dispersion degree of each compartment. a) Abundance of RNA distributed across each compartment. b, c, d, e, f) The number of compartments with which a single compartment can interact in multilabel target data. The plots represent the following RNA types: mRNA, lncRNA, miRNA, snRNA, and snoRNA, respectively.

**Supplementary Figure 6.**
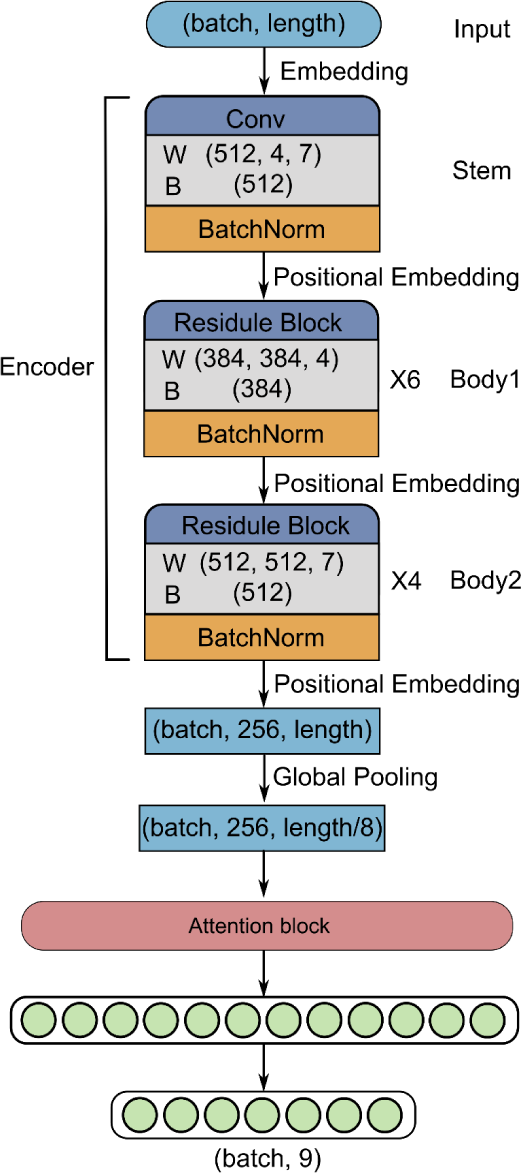
The model architecture of DeepLocRNA.

**Supplementary Figure 7.**
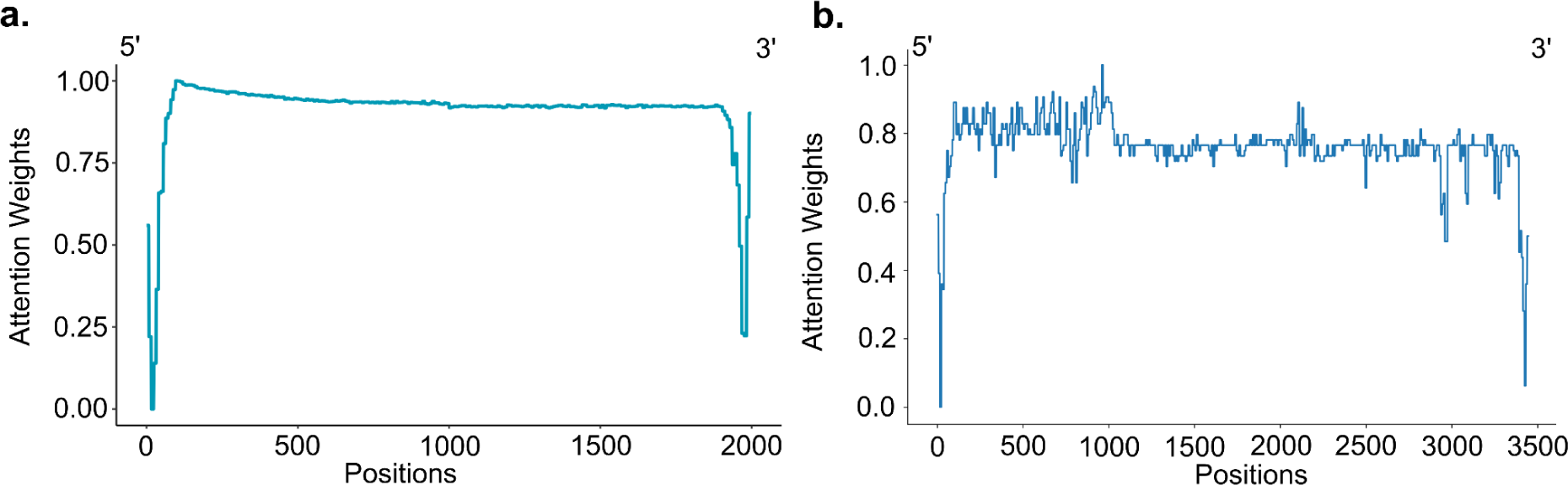
Attention weights analysis across the primary sequences. **a**) This plot displays the attention weights extracted from the attention layer, focusing on the two ends of the sequence. These attention weights provide insights into the neural network’s concentration on specific elements at the beginning and end of the sequence, shedding light on the model’s processing priorities. **b**) The attention weights for the ACTB gene are visualized across its entire sequence. This comprehensive view allows for a detailed examination of how the model allocates attention to different segments of the gene.

**Supplementary Figure 8.**
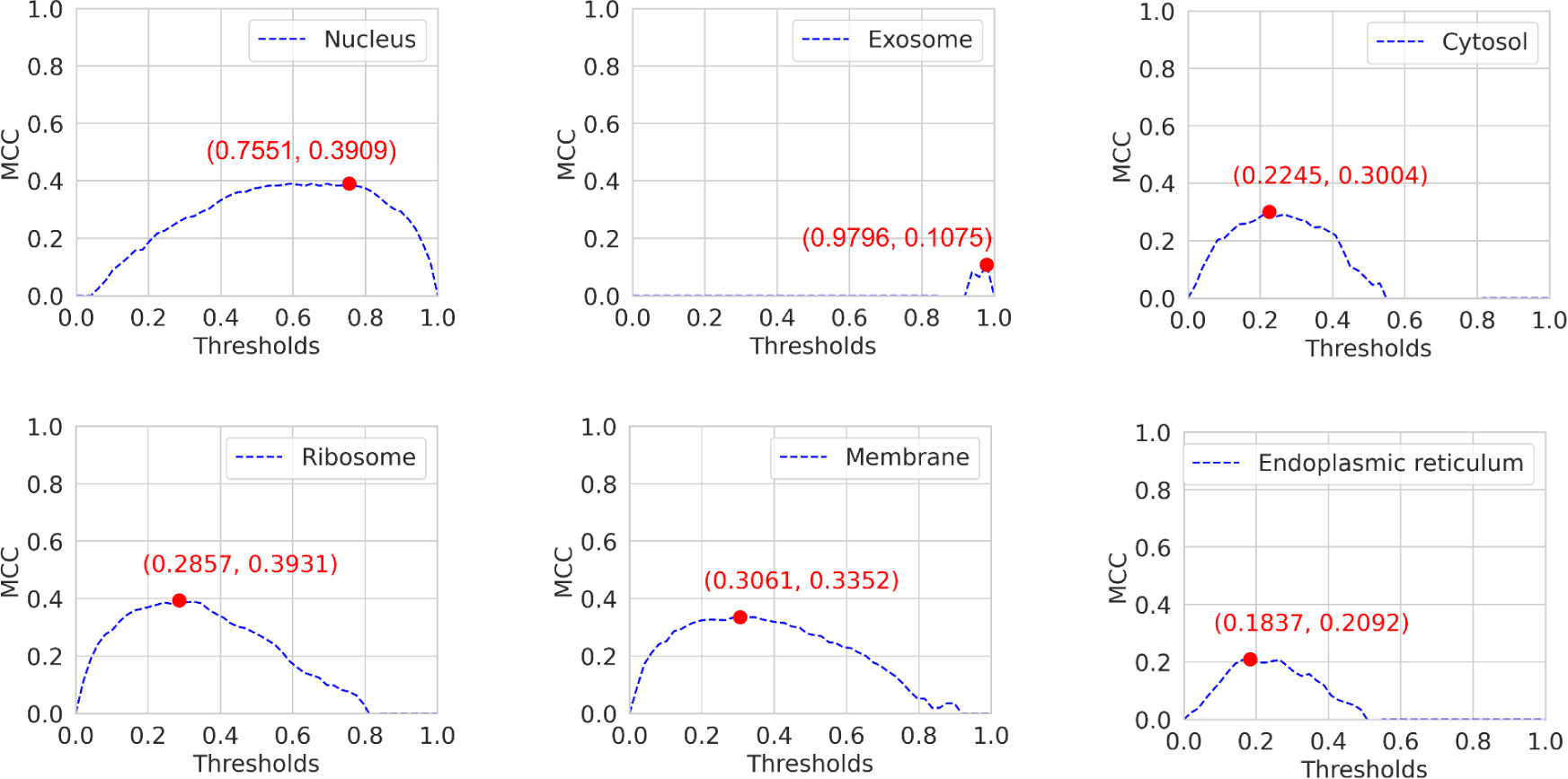
The best MCC score of mRNA across 6 different compartments. The thresholds were initialized by evenly dividing 50 numbers from 0 to 1. The coordinates in red represent the best thresholds and MCC values.

**Supplementary Figure 9.**
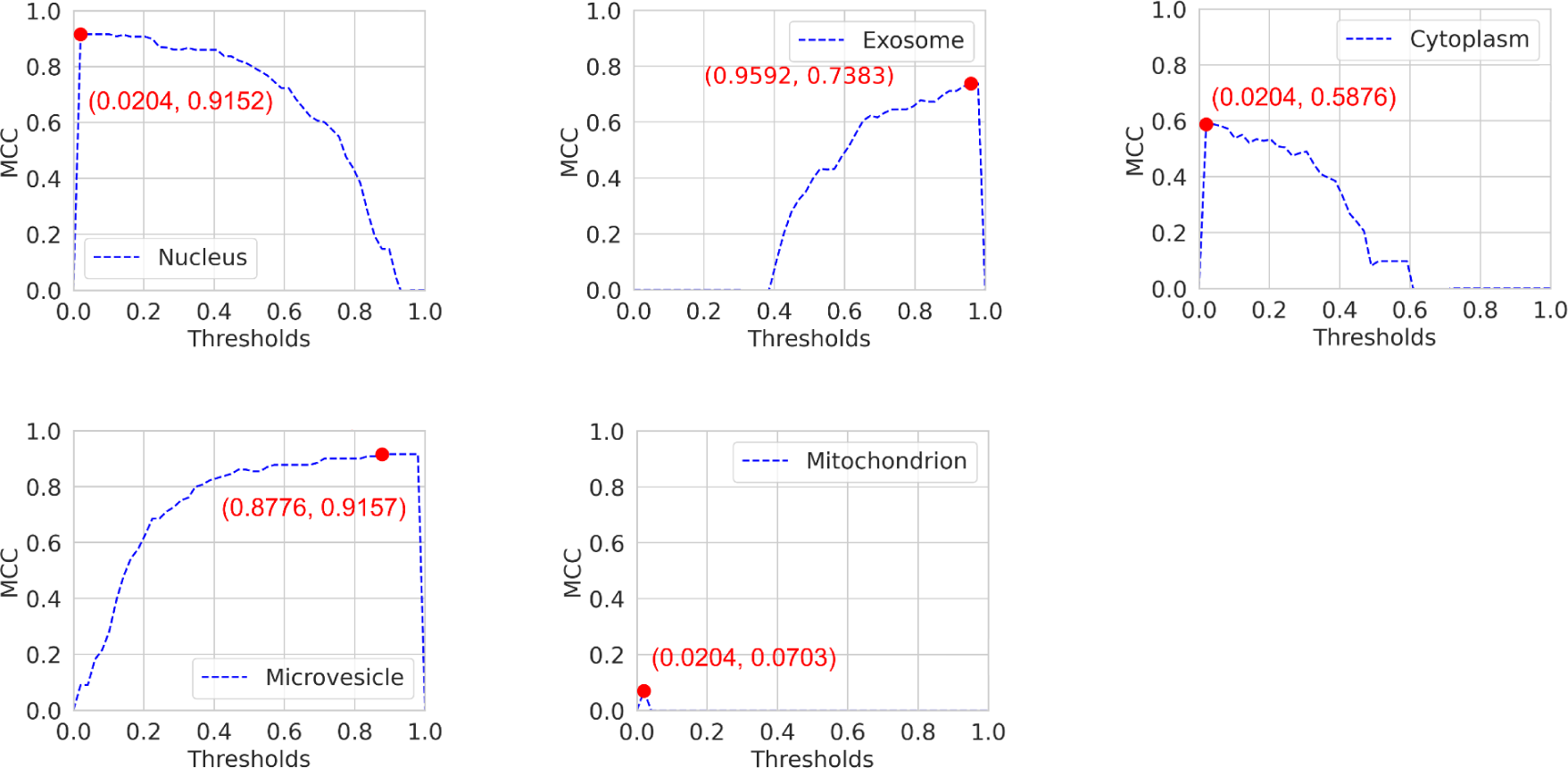
The best MCC score of miRNA across 5 different compartments. The thresholds were initialized by evenly dividing 50 numbers from 0 to 1. The coordinates in red represent the best thresholds and MCC values. Compartments with genes less than 20 were removed, Cytoplasm was shown to replace cytosol because of its rare data.

**Supplementary Figure 10.**
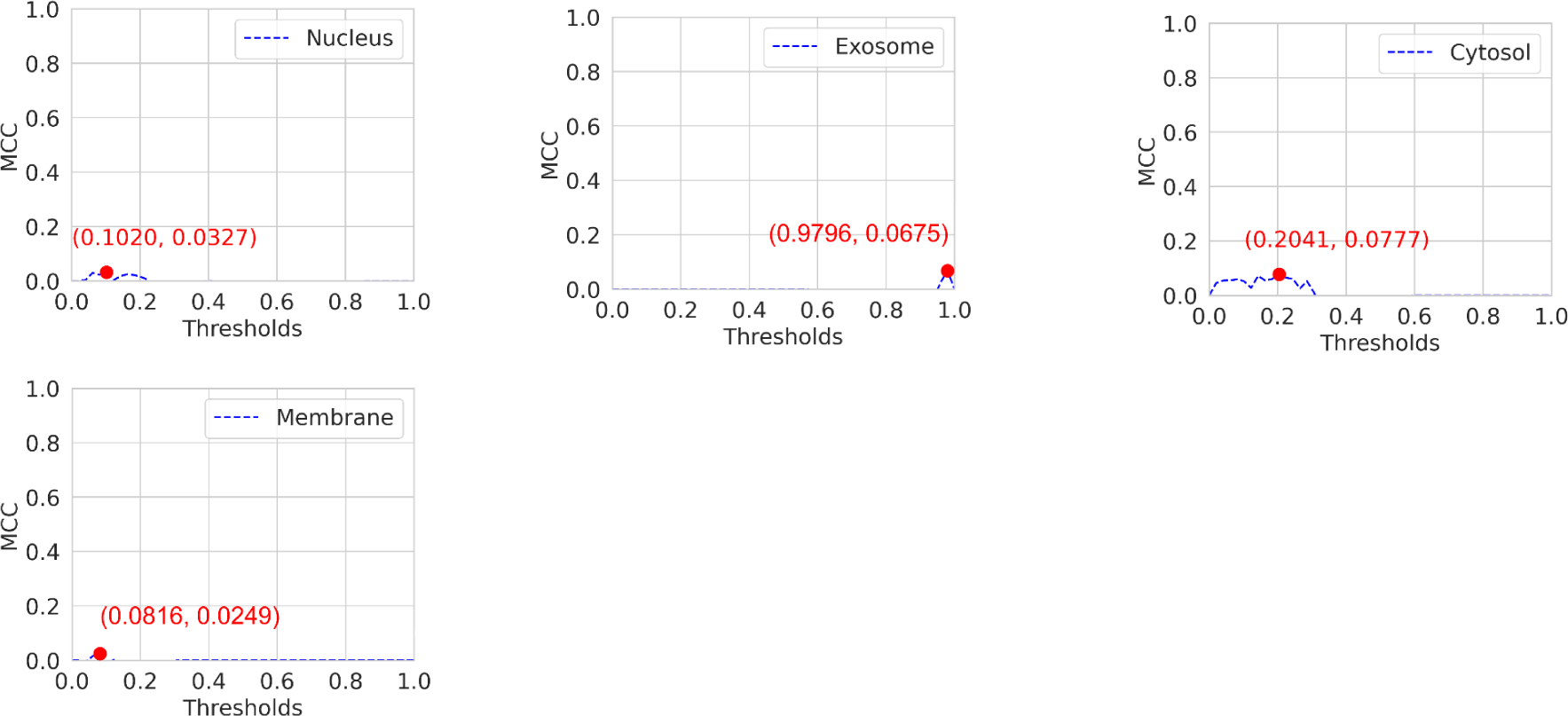
The best MCC score of lncRNA across 4 different compartments. The thresholds were initialized by evenly dividing 50 numbers from 0 to 1. The coordinates in red represent the best thresholds and MCC values. Compartments with genes less than 20 were removed.

**Supplementary Figure 11.**
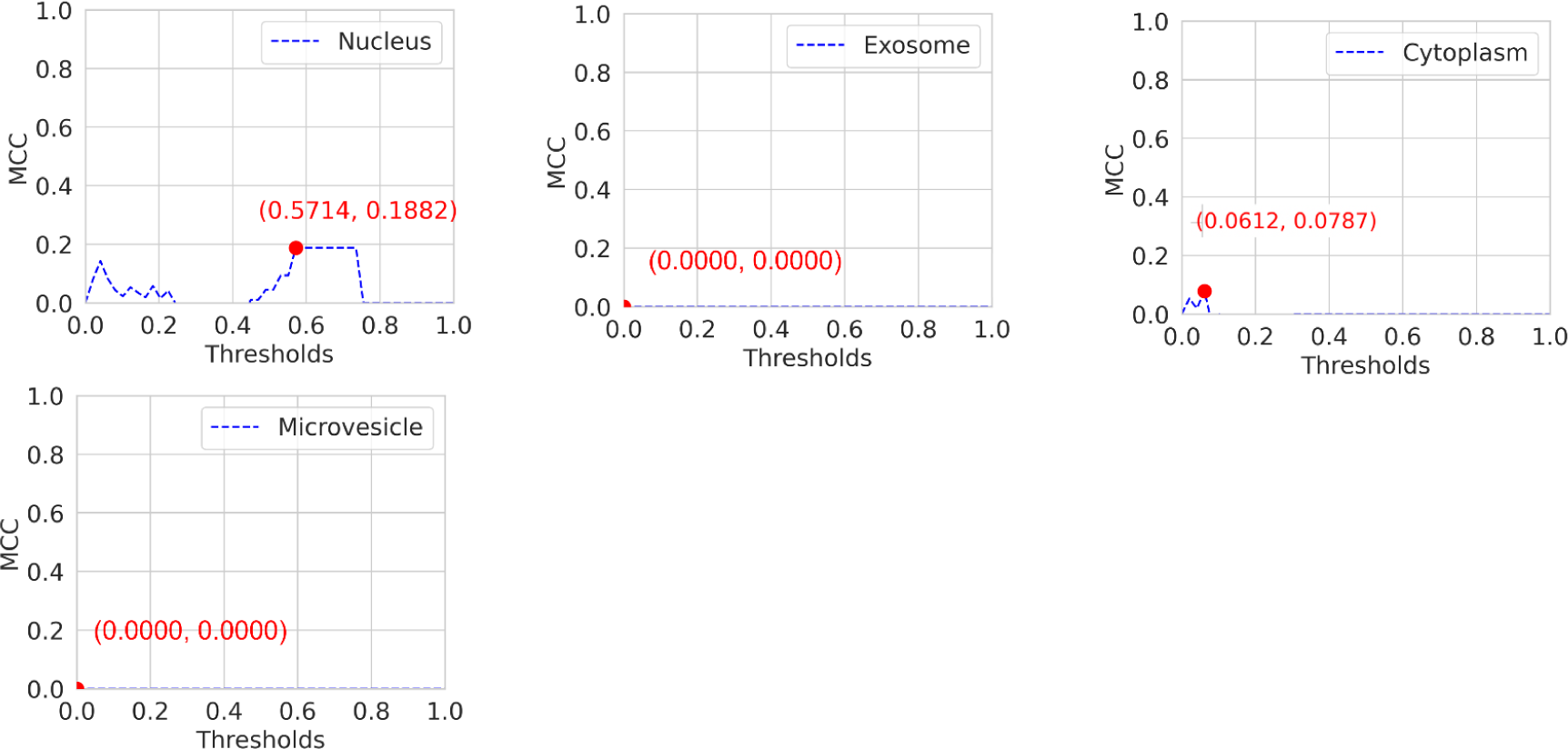
The best MCC score of snoRNA across 4 different compartments. The thresholds were initialized by evenly dividing 50 numbers from 0 to 1. The coordinates in red represent the best thresholds and MCC values. Compartments with genes less than 20 were removed, Cytoplasm was shown to replace cytosol because of its rare data.

**Supplementary Figure 12.**
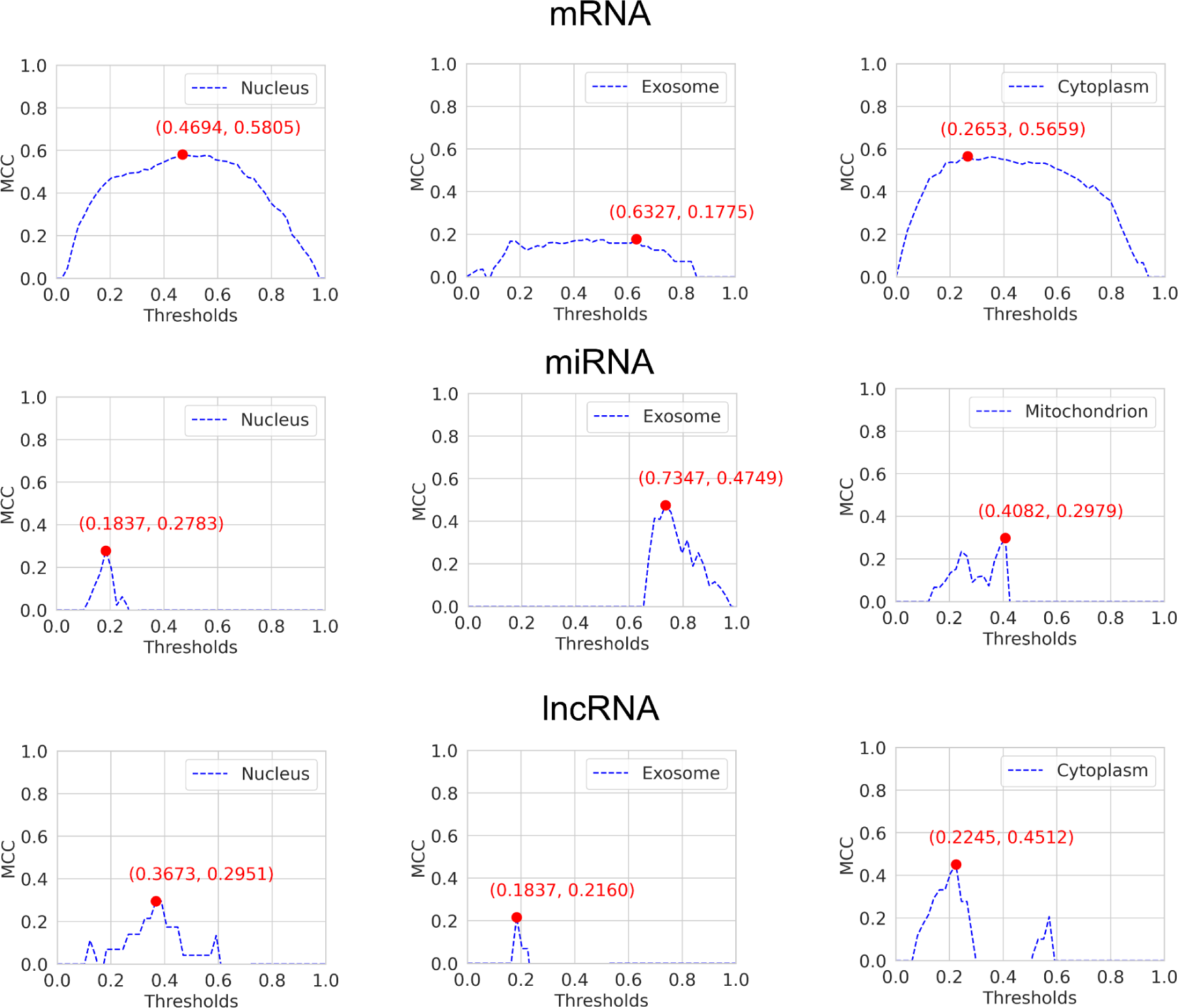
The best MCC score of 3 RNA types in the mouse model. The thresholds were initialized by evenly dividing 50 numbers from 0 to 1. The coordinates in red represent the best thresholds and MCC values. Compartments with genes less than 20 were removed, Cytoplasm was shown to replace cytosol when it has rare data.

**Supplementary Figure 12.**
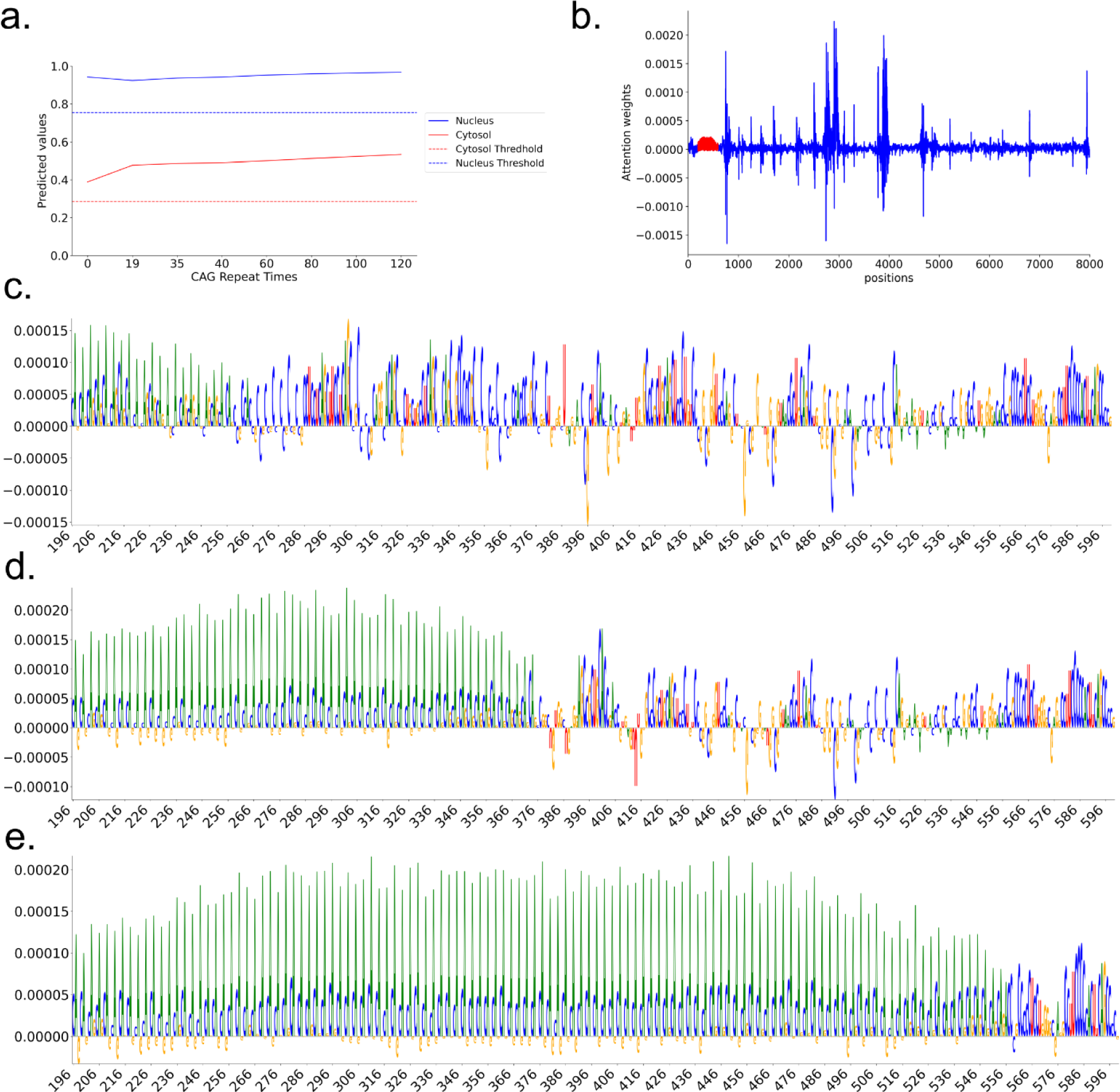
Analysis of the attribution of expanded CAG repeats. a) Variant times of CAG repeats and corresponding prediction probabilities by DeepLocRNA. On the x-axis, ’19’ represents the initial CAG repeat at the 5’ end of the HTT gene, while ’0’ signifies the complete replacement of all CAG repeats in the HTT gene with random nucleotides. b) IG values along the entire HTT gene sequence, with red regions denoting perturbed CAG repeats. c, d, e) Presentation of three levels of expanded CAG repeats and their attribution scores post-mutation, with ’19,’ ’60,’ and ’120’ repeats of CAG shown, respectively.

**Supplementary table 1.** Comparative analysis of DeepRBPLoc and its counterparts using the segregated benchmark dataset. Bold numbers indicate the highest values observed across all comparisons within a given compartment.

| Tools | Compartments | F1 | MCC | AUROC | AUPRC |
| --- | --- | --- | --- | --- | --- |
| DM3Loc | Nucleus | 0.8256 | 0.3516 | 0.7745 | 0.8763 |
|  | Exosome | 0.9959 | 0 | 0.7273 | 0.9964 |
|  | Cytosol | 0.0242 | 0.0625 | 0.7399 | 0.3203 |
|  | Ribosome | **0.4680** | **0.3096** | **0.7592** | **0.5528** |
|  | Membrane | 0.264 | 0.2589 | 0.7546 | **0.4439** |
|  | ER | 0.011 | 0.045 | 0.6980 | 0.2563 |
| iLoc-mRNA | Nucleus | 0 | 0 | 0.5 | 0.651 |
|  | Cytosol | 0.242 | 0.026 | 0.511 | 0.149 |
|  | Ribosome | 0.368 | 0.133 | 0.572 | 0.333 |
|  | ER | 0.123 | 0.017 | 0.516 | 0.104 |
| mRNALoc | Nucleus | 0.391 | 0.1461 | 0.565 | 0.710 |
|  | Cytoplasm | 0.466 | -0.059 | 0.452 | 0.480 |
|  | ER | **0.175** | 0.0621 | 0.546 | 0.095 |
| DeepLocRNA<br>(training from scratch) | Nucleus | 0.7539 | 0.1592 | 0.7429 | 0.8561 |
|  | Exosome | 0.9961 | 0 | 0.7413 | 0.9969 |
|  | Cytosol | 0.0541 | 0.0633 | 0.7404 | **0.3459** |
|  | Ribosome | 0.0692 | 0.0159 | 0.7270 | 0.5209 |
|  | Membrane | 0.0406 | 0.1473 | 0.7446 | 0.4226 |
|  | ER | 0 | 0 | 0.6735 | 0.2303 |
| DeepLocRNA<br>(instructive fine-tuning + weighting) | Nucleus | **0.8341** | 0.3516 | 0.7647 | 0.8681 |
|  | Exosome | **0.9961** | 0 | 0.7537 | 0.9968 |
|  | Cytosol | 0.2016 | **0.1627** | 0.7462 | 0.3383 |
|  | Ribosome | 0.3860 | 0.2460 | 0.7564 | 0.5509 |
|  | Membrane | **0.3001** | 0.2569 | 0.7532 | 0.4337 |
|  | ER | 0.0702 | **0.0474** | **0.7217** | **0.2836** |
| DeepLocRNA<br>(instructive fine-tuning) | Nucleus | 0.8284 | **0.3653** | **0.7760** | **0.8764** |
|  | Exosome | 0.9960 | 0 | **0.7633** | **0.9971** |
|  | Cytosol | **0.0921** | 0.1156 | **0.7476** | 0.3317 |
|  | Ribosome | 0.3382 | 0.2582 | 0.7546 | 0.5475 |
|  | Membrane | **0.2677** | **0.2602** | **0.7551** | 0.4386 |
|  | ER | 0 | 0 | 0.6644 | 0.2295 |

**Supplementary table 2.** Performance of the unified model. The bold numbers represent the larger values when compared with the instructive fine-tuning model (left) and training from scratch model (right). The AUROC of exosome and microvesicle in snoRNA is nan because there are no false negative predictions calculated.

| RNA Species | Compartments | F1 | MCC | AUROC | AUPRC |
| --- | --- | --- | --- | --- | --- |
| mRNA | Nucleus | **0.8252** \| 0.8019 | **0.3840** \| 0.3611 | **0.7752** \| 0.7654 | **0.8761** \| 0.8700 |
|  | Exosome | **0.9961** \| 0.9960 | 0 \| 0 | **0.7598** \| 0.7325 | **0.9968** \| 0.9960 |
|  | Cytosol | **0.0109** \| 0 | **0.0270** \| 0.0024 | **0.7435** \| 0.7169 | **0.3244** \| 0.2943 |
|  | Ribosome | **0.3489** \| 0.2776 | **0.2678** \| 0.2101 | **0.7607** \| 0.7539 | **0.5624** \| 0.5381 |
|  | Membrane | **0.2627** \| 0.0107 | **0.2560** \| 0.0532 | **0.7551** \| 0.7416 | **0.4412** \| 0.4164 |
|  | ER | **0.0039** \| 0 | **0.0137** \| 0 | **0.6655** \| 0.5966 | **0.2232** \| 0.1600 |
|  | **Micro** | **0.7703** \| 0.7353 | / | **0.9281** \| 0.9229 | **0.8838** \| 0.8771 |
| miRNA | Nucleus | **0.8872** \| 0.7973 | **0.8524** \| 0.7499 | **0.9760** \| 0.9028 | **0.8987** \| 0.7988 |
|  | Exosome | 0.9149 \| **0.9431** | 0.2865 \| **0.4801** | 0.9327 \| **0.9526** | 0.9879 \| **0.9938** |
|  | Cytoplasm | **0.0762** \| 0 | **0.1057** \| 0.0056 | **0.9008** \| 0.8008 | **0.4808** \| 0.3750 |
|  | Microvesicle | 0.9633 \| **0.9827** | 0.8470 \| **0.9056** | 0.9757 \| **0.9776** | 0.9932 \| **0.9938** |
|  | Mitochondrion | 0 \| 0 | 0 \| 0 | 0.5556 \| **0.7018** | 0.0241 \| **0.2431** |
|  | **Micro** | **0.8970** \| 0.8874 | / | **0.9909** \| 0.9803 | **0.9726** \| 0.9562 |
| lncRNA | Nucleus | **0.0965** \| 0.0223 | 0.0174 \| **0.0477** | 0.5354 \| **0.5483** | 0.2701 \| **0.2724** |
|  | Exosome | 0.9864 \| **0.9867** | **0.0019** \| 0 | 0.6582 \| **0.6801** | 0.9863 \| **0.9872** |
|  | Cytosol | 0 \| 0 | 0 \| 0 | **0.5872** \| 0.5723 | **0.1021** \| 0.0918 |
|  | Membrane | 0 \| 0 | **0.0015** \| 0 | 0.5520 \| **0.5645** | 0.0461 \| **0.0483** |
|  | **Micro** | **0.8273** \| 0.8213 | / | **0.9666** \| 0.9663 | 0.8859 \| **0.8911** |
| snoRNA | Nucleus | **0.1182** \| 0.0161 | **0.0993** \| 0.0437 | **0.6541** \| 0.6340 | **0.3320** \| 0.2868 |
|  | Exosome | 1 \| 1 | 0 \| 0 | // | // |
|  | Cytoplasm | 0 \| 0 | 0 \| 0 | **0.6363** \| 0.5346 | **0.1878** \| 0.1075 |
|  | Microvesicle | **0.9991** \| 0.9988 | 0 \| 0 | // | // |
|  | **Micro** | **0.9333** \| 0.9290 | / | 0.9902 \| 0.9898 | **0.9793** \| 0.9777 |
| Overall | Nucleus | **0.7955** \| 0.7708 | **0.5085** \| 0.5050 | **0.8306** \| 0.8298 | **0.8588** \| 0.8483 |
|  | Exosome | 0.9877 \| **0.9881** | 0.2577 \| **0.4049** | 0.9024 \| **0.9040** | 0.9961 \| **0.9964** |
|  | Cytosol | **0.0097** \| 0 | **0.0269** \| 0.0018 | **0.7650** \| 0.7547 | **0.3043** \| 0.2660 |
|  | Ribosome | **0.3489** \| 0.2776 | **0.3060** \| 0.2473 | **0.8394** \| 0.8388 | **0.5622** \| 0.5356 |
|  | Membrane | **0.2559** \| 0.0103 | **0.2676** \| 0.0548 | 0.7591 \| **0.8050** | **0.4323** \| 0.4017 |
|  | ER | **0.0039** \| 0 | **0.0142** \| 0 | 0.7107 \| **0.7189** | **0.2232** \| 0.1556 |
|  | Microvesicle | 0.9703 \| **0.9836** | 0.9674 \| **0.9812** | **0.9995** \| 0.9994 | **0.9952** \| 0.9937 |
|  | Mitochondrion | 0 \| 0 | 0 \| 0 | 0.9599 \| **0.9638** | 0.0241 \| **0.2241** |
|  | **Micro** | **0.7831** \| 0.7626 | / | **0.9445** \| 0.9413 | **0.8957** \| 0.8922 |

**Supplementary table 3.**
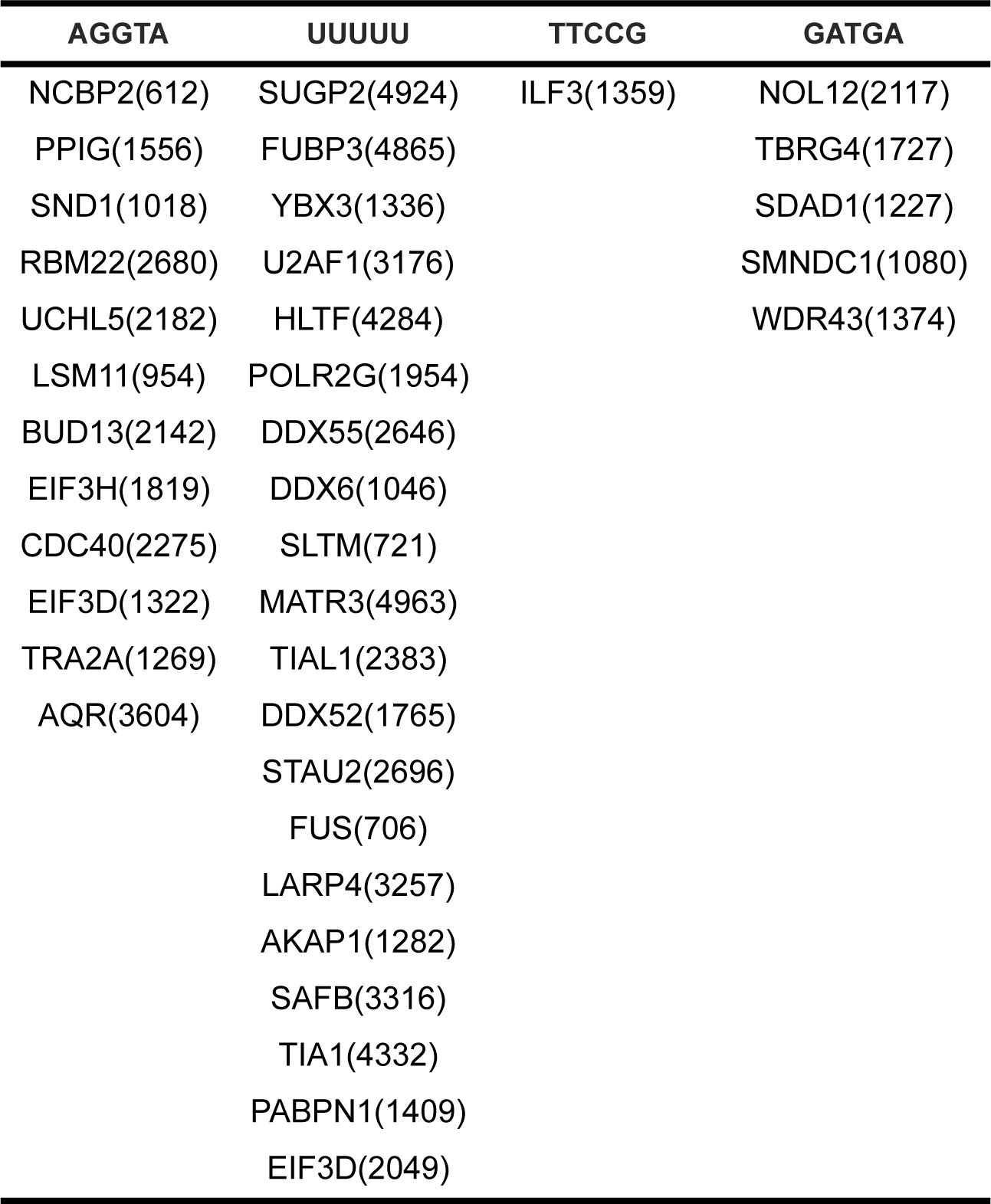
4 extracted motifs that were also found in RBPnet. The proteins displayed following the motifs are those that bind to these specific motifs. The numbers in parentheses represent the support count of these motifs in RNA-binding proteins (RBPs).

| AGGTA | UUUUU | TTCCG | GATGA |
| --- | --- | --- | --- |
| NCBP2(612) | SUGP2(4924) | ILF3(1359) | NOL12(2117) |
| PPIG(1556) | FUBP3(4865) |  | TBRG4(1727) |
| SND1(1018) | YBX3(1336) |  | SDAD1(1227) |
| RBM22(2680) | U2AF1(3176) |  | SMNDC1(1080) |
| UCHL5(2182) | HLTF(4284) |  | WDR43(1374) |
| LSM11(954) | POLR2G(1954) |  |  |
| BUD13(2142) | DDX55(2646) |  |  |
| EIF3H(1819) | DDX6(1046) |  |  |
| CDC40(2275) | SLTM(721) |  |  |
| EIF3D(1322) | MATR3(4963) |  |  |
| TRA2A(1269) | TIAL1(2383) |  |  |
| AQR(3604) | DDX52(1765) |  |  |
|  | STAU2(2696) |  |  |
|  | FUS(706) |  |  |
|  | LARP4(3257) |  |  |
|  | AKAP1(1282) |  |  |
|  | SAFB(3316) |  |  |
|  | TIA1(4332) |  |  |
|  | PABPN1(1409) |  |  |
|  | EIF3D(2049) |  |  |

**Supplementary table 4.** Summary of the number of RNAs across nine subcellular compartments of human and mouse.

| Species | Compartments | Nucleus | Microvesicle | Mitochondrion | Exosome | Cytosol | Cytoplasm | Ribosome | Membrane | ER |
| --- | --- | --- | --- | --- | --- | --- | --- | --- | --- | --- |
| Human | lncRNA | 836 | 1 | 1 | 3092 | 246 | 103 | 18 | 124 | 2 |
|  | miRNA | 472 | 1466 | 32 | 1605 | 0 | 234 | 1 | 0 | 0 |
|  | snRNA | 13 | 16 | 0 | 19 | 0 | 3 | 0 | 0 | 0 |
|  | snoRNA | 115 | 475 | 0 | 483 | 4 | 40 | 0 | 1 | 0 |
|  | mRNA | 11916 | 0 | 0 | 17136 | 2337 | 9646 | 5207 | 3231 | 1975 |
|  | **Sum** | **13352** | **1958** | **33** | **22335** | **2587** | **10026** | **5226** | **3356** | **1977** |
| Mouse | lncRNA | 36 | 0 | 0 | 33 | 2 | 22 | 0 | 0 | 0 |
|  | miRNA | 62 | 8 | 95 | 301 | 0 | 7 | 0 | 0 | 0 |
|  | mRNA | 2119 | 0 | 1 | 782 | 11 | 1491 | 2 | 1 | 8 |
|  | **Sum** | **2271** | **8** | **96** | **1116** | **13** | **1520** | **2** | **1** | **8** |

